# Domain-Specific Functional Network Adaptations Supporting Dual-Task Performance in Older Adults

**DOI:** 10.1101/2025.09.28.678786

**Authors:** Yan Deng, AmirHussein Abdolalizadeh, Kayson Fakhar, Tina Schmitt, Karsten Witt, Carsten Gießing, Jochem W. Rieger, Christiane M. Thiel

**Affiliations:** Biological Psychology Lab, Department of Psychology, School of Medicine and Health Sciences, Carl von Ossietzky Universität Oldenburg, Oldenburg, Germany; MRC Cognition and Brain Sciences, University of Cambridge, Cambridge, United Kingdom; Institute of Computational Neuroscience, University Medical Center Hamburg Eppendorf, Hamburg, Germany; Neuroimaging Unit, School of Medicine and Health Sciences, Carl von Ossietzky Universität Oldenburg, Oldenburg, Germany; Department of Neurology, School of Medicine and Health Sciences, Carl von Ossietzky Universität Oldenburg, Oldenburg, Germany; Research Center Neurosensory Science, Carl von Ossietzky Universität Oldenburg, Oldenburg, Germany; Applied Neurocognitive Psychology Lab, Department of Psychology, School of Medicine and Health Sciences, Carl von Ossietzky Universität Oldenburg, Oldenburg, Germany

**Keywords:** task-based functional connectivity, cognitive-motor dual-task, brain-behavior association, domain-specific network, regularized generalized canonical correlation analysis (RGCCA)

## Abstract

Aging is associated with declines in both motor and cognitive functions, which are well captured by dual-task gait paradigms. However, the functional brain network mechanisms supporting motor and cognitive aspects of dual-task performance in aging remain unclear. We examined 40 older adults (50-80 years) and 20 younger adults (20-40 years) who performed a motor single-task (pedaling), a cognitive single-task (Go/NoGo), and a combined cognitive–motor dual-task during functional magnetic resonance Imaging (fMRI) using a custom-built MRI-compatible pedaling device. Behaviorally, older adults showed significant dual-task costs in motor performance, while cognitive performance was preserved. Neurally, older adults showed selective increases in connectivity within executive and motor-planning regions of cognitive networks, consistent with compensatory recruitment, whereas motor networks underwent broader reorganization, with strengthened frontoparietal control circuits but weakened cerebello-parietal and sensorimotor pathways. Multivariate analyses further revealed age-related differences in latent connectivity– behavior relationships: motor-network patterns in older adults were more dispersed, reflecting heterogeneous reorganization, whereas cognitive-network patterns were more overlapping across groups, suggesting relative preservation. These findings suggest that aging involves a domain-specific balance of resilience and vulnerability across brain networks and highlight motor-network adaption as a promising target for understanding why some older adults maintain function while others decline.

## 1. Introduction

Aging is associated with a gradual decline in both cognitive and motor functions (Deary et al., 2009; Niederer et al., 2021; Wilson et al., 2020), which can adversely impact daily activities in older adults. One informative approach for evaluating these changes is the cognitive-motor dual-task paradigm, in which individuals perform a motor task (e.g., walking) while simultaneously engaging in a cognitive activity (e.g., talking or serial subtraction). This paradigm has proven effective for detecting early signs of cognitive impairment, including mild cognitive impairment (MCI) (Ali et al., 2024; Bishnoi & Hernandez, 2021; Yang et al., 2020), dementia (Collyer et al., 2022; Montero-Odasso et al., 2018), Alzheimer’s disease (Belghali et al., 2017; Longhurst et al., 2023), and Parkinson’s disease (Kim et al., 2024; Longhurst et al., 2023). Early identification of these deficits, along with understanding their neural correlates, is critical for developing timely interventions and potentially slowing the progression of neurodegenerative conditions.

In healthy aging, findings on dual-task performance are inconsistent. Some studies report slower, more variable gait and greater cognitive-motor interference compared with younger adults (Hollman et al., 2007a; Lino et al., 2023), whereas others show domain-specific declines—for example, preserved cognitive performance but impaired motor control, such as increased gait variability (Bürki et al., 2017) or altered posture (Bohle et al., 2019). In contrast, St George et al. (2022) observed smaller walking-speed reductions in older than younger adults, yet greater cognitive performance declines. To capture these diverse patterns, Collyer et al. (2022) proposed classifying older adults into four groups: (1) dual decline in gait and cognition, (2) gait decline only, (3) cognitive decline only, and (4) non-decliners, highlighting the behavioral heterogeneity of dual-task effects. Linking such behavioral variability to underlying neural mechanisms requires imaging methods that can separate motor- and cognitive-related processes during dual-task performance.

Most prior neuroimaging studies of dual-tasking in older adults have relied on functional near-infrared spectroscopy (fNIRS) during walking paradigms (Holtzer et al., 2011; St George et al., 2022; Udina et al., 2021). While informative, fNIRS is restricted to superficial cortical regions, limiting insights into subcortical motor structures. To overcome this limitation, MRI-compatible pedaling tasks have been developed, offering a gait-like motor activity that minimizes head motion while allowing whole-brain measurement (Bürki et al., 2017; Papegaaij et al., 2017; Promjunyakul et al., 2015; Reinhardt et al., 2020). These studies demonstrated that pedaling is a suitable proxy for gait, engaging motor and cerebellar regions, and that dual-tasking modulates both motor and prefrontal activity. However, findings remain inconsistent across studies, with dual-task interference linked to under-, over-, or mixed activation patterns (Leone et al., 2017).

To move beyond regional activation and capture coordinated network dynamics, recent work has turned to functional connectivity approaches. Such a network-level perspective is well suited to dual-task research, because it assesses how motor, cognitive, and control systems interact dynamically under varying task demands (Vinehout et al., 2019). Yet task-based functional connectivity studies of gait-like cognitive-motor dual-task in aging remain scarce, especially those incorporating gait-like movements. Most prior work was limited to resting-state analyses, which relied on relating dual-task performance outside the scanner to changes in resting-state functional connectivity (Boyne et al., 2018; Droby et al., 2022; Yuan et al., 2015).

The present study addresses this gap by examining task-based functional connectivity during cognitive-motor dual-task in older and younger adults using an MRI-compatible pedaling paradigm. By contrasting dual-task with single-task conditions, we isolated motor-specific and cognitive-specific networks, enabling us to assess domain-specific neural reorganization. To link connectivity with performance, we employed RGCCA, a multivariate integration approach (Rohart et al., 2017; Tenenhaus et al., 2017). This framework allowed us to identify broad multivariate associations between functional connectivity and dual-task behavior, to highlight specific pairwise relationships, and to descriptively illustrate age-related variation in these associations.

Our study aimed to (i) characterize domain-specific differences in task-based functional connectivity comparing older adults with younger adults, (ii) assess how these age-related connectivity patterns relate to dual-task performance, and (iii) illustrate how these associations vary with age across task domains. This approach provides new insights into how the aging brain dynamically adapts functional networks to balance resilience and vulnerability under cognitive–motor demands.

## 2. Methods

### 2.1 Participants

We recruited 62 healthy, right-handed participants through the University of Oldenburg campus bulletin board and local advertisements. Participants were recruited into two age groups: older adults (n = 40; 19 males), aged 50–80 years (mean = 67.6, SD = 7.09), and young adults (n = 20; 11 males), aged 20–40 years (mean = 28.0, SD = 4.87). All participants reported no history of neurological, psychiatric, or motor-related disorders. Demographic information, including age, gender, educational level, and medical history, was obtained through a self-report questionnaire.

The study protocol was approved by the research ethics committee of the University of Oldenburg and conducted in accordance with the declaration of Helsinki (Drs.EK/2020/062-02). Written informed consent was obtained from all participants. Two participants from the older group were excluded due to excessive head motion that induced stripe artefacts in the fMRI images.

### 2.2 Cognitive-motor dual-task

#### Experimental design

The fMRI paradigm included three task conditions: a cognitive single-task, a motor single-task, and a combined cognitive-motor dual-task (Fig. 1). The order of the two single-task runs was counterbalanced across participants; the dual-task run was always presented last. Each task began with a 10-s instruction period, followed by eight cycles of alternating task blocks. Each block lasted 25.8 s and was followed by a 10-s rest, resulting in a total duration of 296.4 s (≈ 4.94 min) per task. Participants completed two practice sessions: (i) a computer-based session outside the scanner to ensure task comprehension, and (ii) an in-scanner familiarization to practice in the supine position under MRI conditions. Both practice sessions included two blocks of each task, each lasted about 1.5 min. All three tasks were presented and responses recorded using Presentation (version 22.1, Neurobehavioral System, Inc).

**Figure 1.**
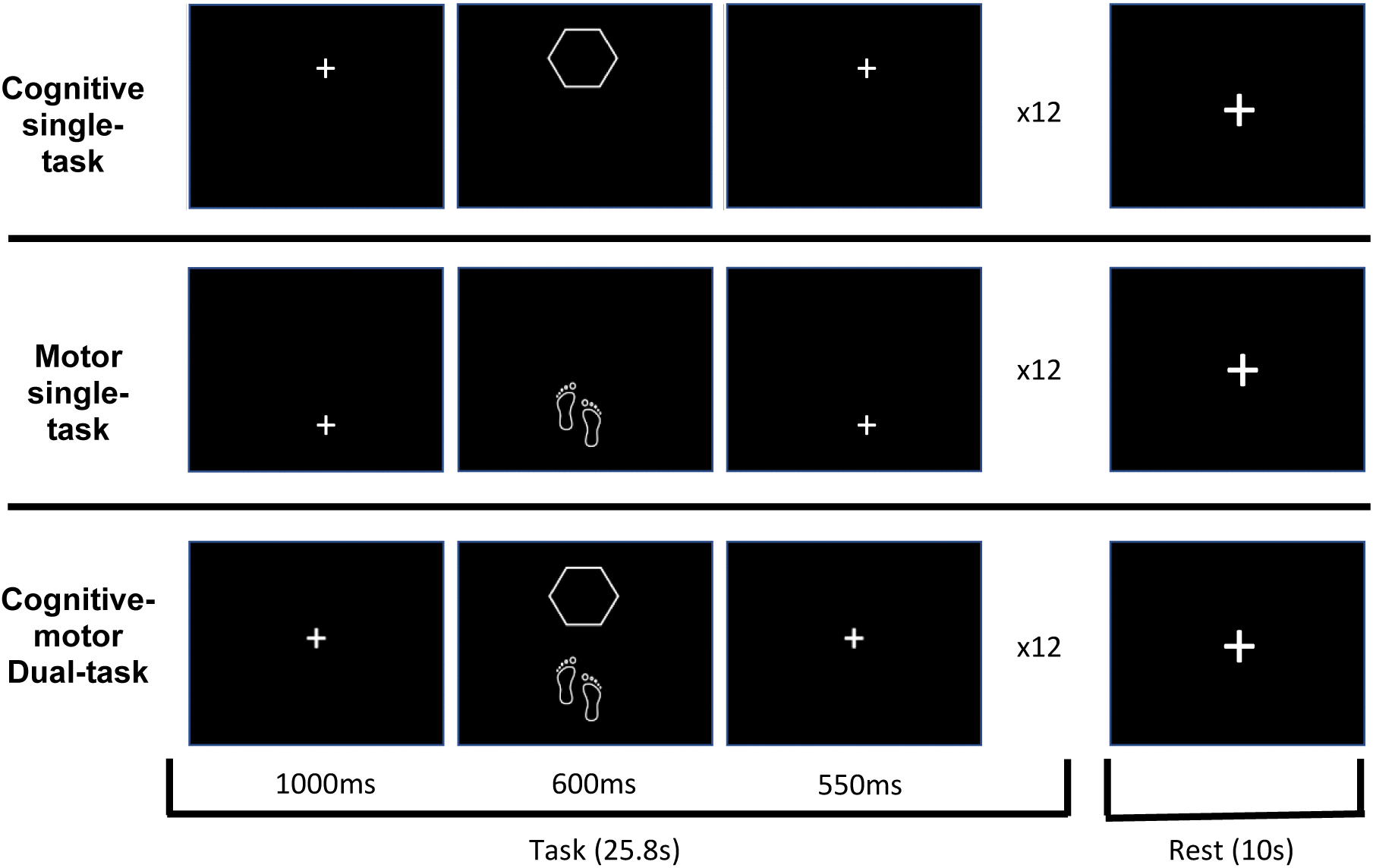
Experimental paradigm: illustration of a single trial from each task condition. Upper row: cognitive single-task (Go trial requiring button press; *“single cognitive go”*). Middle row: motor single-task (foot symbol requiring pedal press; *“single motor”*). Lower row: combined cognitive–motor dual-task (Go trial with a concurrent button and pedal press; *“dual go”*). Each task block consisted of 12 trials (75% Go, 25% NoGo; NoGo trials not shown), followed by a rest block.

#### MRI-compatible pedal device

Custom-made, MR-compatible foot pedals (Fig. 2A) were used to elicit alternating dorsiflexion and plantarflexion movements, simulating gait-related activity in a supine position. The pedals, constructed from polyvinyl chloride (PVC), were securely mounted on a base plate and fixed within the scanner bed rails to ensure stable positioning throughout the measurement period (Fig. 2B). Pedal movements generated signals that were transmitted via optical fibres to an interface unit outside the scanner room, where they were converted into electrical signals and transferred via USB connection to a PC. The output included a numerical string representing the real-time position of each pedal along with a timestamp, recorded with a temporal resolution of 0.1 ms. This setup enabled precise monitoring of foot movements, allowing accurate assessment of motor performance during the experiment.

**Figure 2.**
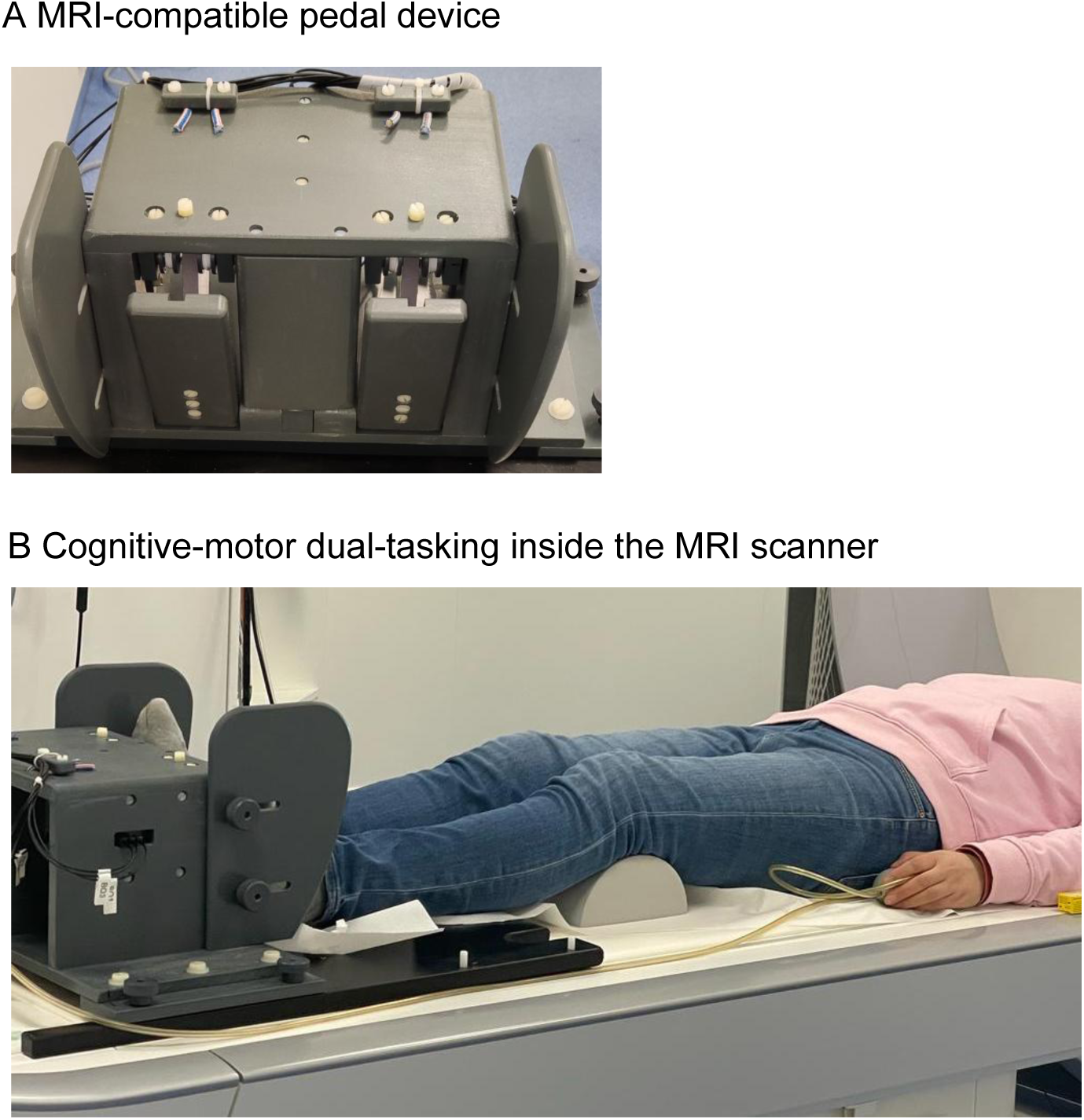
(A) MRI-compatible pedal device. (B) Pedals mounted securely on a base plate and fixed within the scanner bed rails to ensure stable positioning throughout the cognitive-motor dual-tasking measurements. Participants were comfortably positioned on the scanner bed and the pedal device was adjusted to their height to allow natural leg extension and optimal foot movements. (Note: The photograph includes one of the authors, who has provided consent for its publication.)

#### Cognitive single-task

The cognitive single-task employed a Go/NoGo inhibitory control paradigm, adapted from the Human Connectome Project (HCP) in Aging (Bookheimer et al., 2019). Participants were instructed to press a button with their right index finger as quickly as possible in response to various white-outlined geometric shapes presented on a black background, and to withhold their response to two specific shapes. Six shapes—plaques, trapezoids, pentagons, hexagons, octagons, and parallelograms— served as Go stimuli (*“*single cognitive Go”), while squares and circles served as NoGo stimuli (“single cognitive NoGo”). Each shape was presented for 600 ms in the upper half of the screen, with stimuli presented in randomized order. Each task block consisted of 12 trials and was followed by a 10-s rest period, during which a larger fixation cross was displayed. The task was repeated eight times, resulting in a total task duration of 296.4 s (≈ 4.94 min). In total, the task comprised 96 trials, with 75% Go trials and 25% NoGo trials.

#### Motor single-task

In the motor single-task, a foot symbol was presented on the screen in the lower half of the screen at a fixed rhythm to simulate a walking cadence. Participants were instructed to press one pedal with a foot in response to each symbol and to alternate between feet across trials (“single motor”). Pedal signals were recorded during the stimulus presentation and for an additional 550 ms, capturing the full movement cycle from pedal press to release. Each stepping cycle lasted 2.15 seconds, comprising a 1-s fixation period (interstimulus interval), 600 ms of stimulus presentation, and 550 ms of additional pedal signal recording. Each cycle was repeated 12 times per block. As with the cognitive task, each task block was repeated eight times and followed by a 10-s rest period, during which a larger fixation cross was presented. In total, the motor single-task comprised 96 trials and lasted 296.4 s (≈ 4.94 min).

#### Cognitive-motor dual-task

In the cognitive-motor dual-task, both the geometrical shape and the foot symbol were presented simultaneously on the screen. The shapes appeared in the upper part and the foot symbol in the lower part. Participants were instructed to respond to Go trials by pressing a button with their index finger while simultaneously stepping with one-foot, alternating sides across trials (“Dual Go”). In NoGo trials, they were to withhold the button press while continuing the stepping movement (“Dual NoGo”). Each dual-task trial lasted 2.15 s comprising a 1-s fixation period (interstimulus interval), 600 ms stimulus presentation, and 550 ms of additional pedal signal collection. Each task included 12 trials and was followed by a 10-s rest, during which a larger fixation cross was presented. The task was repeated eight times resulting in a total task duration of 296.4 s (≈ 4.94 min).

### 2.3 Test of Attentional Performance (TAP-M) battery

After completing the dual-task paradigm, participants also performed the Test of Attentional Performance (TAP-M; Zimmermann & Fimm, 2002) to provide a general assessment of attentional and executive functions. Four subtests were administered: (1) The distractibility task assessed the ability to maintain central focus while ignoring unpredictable peripheral stimuli. The outcome measure was the median reaction times (RTs) to central targets in the presence of distractors; (2) The executive control task probed selective attention, inhibition, and cognitive flexibility through responses to specific color–letter or color–number combinations. Outcome was median RTs for correct trials; (3) The divided attention task required simultaneous monitoring of visual and auditory streams. Outcome measures included median RTs for correct detections in each modality, (4) The Go/NoGo task assessed response inhibition. Outcome was median RTs for correct “Go” responses.

### 2.4 Behavioral data analysis

All behavioral data were processed using MATLAB (version 2022b). Behavioral outcomes included mean RT, RT variability (gauged by standard deviations, SD), dual-task cost, and error rate. Cognitive performance was assessed by computing the mean RTs and SDs across all correct Go trials, again separately for single- and dual-task conditions. RT was defined as the time between presentation of the geometric shape and the button press. False responses to NoGo stimuli were counted as errors in cognitive performance, calculated in terms of the number of incorrected responses divided by the total number of NoGo stimuli and reported as error rate in percentage. Motor performance was gauged using mean RTs and SDs, calculated across all pedal presses for each participant under both single- and dual-task conditions. Missed pedal presses were counted as errors in motor performance (missed responses relative to the total number of expected pedal presses, reported as error rate in percentage). RT in motor performance was defined as the time between presentation of the foot symbol and the onset of the pedal displacement. Dual-task costs for both cognitive and motor performance (based on both RTs and SDs) were computed as the relative change between dual-task and single-task performance, expressed as a percentage of single-task performance.

Statistical analyses focused on mean RTs and RT variability using three-way mixed-design analysis of variance (ANOVA) to assess the effects of age (younger vs. older adults; between-subjects), task condition (single vs. dual; within-subjects) and domain (motor vs. cognitive; within-subjects), as well as their interactions. Given the unequal group sizes (40 older adults vs. 20 younger adults), Type III sum of squares was employed, as it provides unbiased estimates in unbalanced designs. Statistical significance was assessed using F-tests (α = 0.05). Effect sizes were reported as partial eta-squared. Analyses were performed in R (v4.4.3) using *afex* package.

In addition, to illustrate age-related performance differences across all behavioral measures, we conducted independent-samples t-tests comparing younger and older adults in each task condition. To account for multiple comparisons, we applied a Bonferroni correction to adjust the significance threshold for multiple testing (16 comparisons), resulting in an adjusted alpha level of *p* < 0.0031 (0.05 /16).

### 2.5 fMRI data acquisition

MRI data were acquired using a 3T Siemens MAGNETOM Prisma scanner (Siemens Healthcare, Erlangen, Germany) equipped with a 64-channel head coil. Participants were positioned comfortably on the scanner bed and the pedal device was adjusted according to their height to allow natural leg extension and effective foot movements (Fig. 2B). To minimize head movement during pedaling, participants were instructed to keep their heads as still as possible. Soft foam pads were used to stabilize the head, and a fixing plaster (Leukosilk) was placed on the participant’s forehead and the boarders of the head coil as a tactile aid, allowing them to detect and self-regulate any unintended head movements. The fMRI session began with a short training phase in which participants performed both the single and dual tasks which lasted approximately 4.5 minutes. After training, task performance was evaluated with regard to alternating pedaling and head motion, and participants received brief feedback before proceeding to the main task which lasted approximately 15 minutes.

A T2*-weighted gradient echo planar imaging (EPI) with BOLD contrast was acquired using the multiband sequence developed by the Center for Magnetic Resonance Research (CMRR), University of Minnesota (Feinberg et al., 2010; Moeller et al., 2010; Xu et al., 2013). The imaging parameters were as follows: repetition time (TR) = 850 ms, echo time (TE) = 30 ms, flip angle = 62°, distance factor = 0%, slice thickness = 2.5 mm, field of view = 192 × 192 mm, matrix size = 76 × 76 voxels, voxel size = 2.5 × 2.5 × 2.5 mm³, 48 slices, multiband factor = 4, acquisition time = 15:26 minutes and 1090 volumes. After completing the task, a high-resolution three-dimensional T1-weighted structural image was acquired for each participant using the following parameters: TR = 2000 ms, TE = 2.07 ms, flip angle = 9°, voxel size = 0.75 × 0.75 × 0.75 mm³, GRAPPA acceleration factor = 2, FOV = 240 × 240 mm, 224 sagittal slices, acquisition time = 6:16 minutes. Finally, a resting-state fMRI was performed using the same 2D multiband EPI sequence with BOLD contrast (TR = 850 ms, TE = 30 ms, flip angle = 62°, slice thickness = 2.5 mm, field of view = 192 x 192 mm, matrix = 76 x 76, voxel size = 2.5 x 2.5 x 2.5 mm³, 48 slices, multiband factor = 4, and 750 volumes).

### 2.6 fMRI data analysis

fMRI data analysis consisted of three steps. First, a task-based voxel-wise GLM analysis (“brain activation analysis”) was performed to identify regions specifically associated with the cognitive and motor demands of dual-task performance (Dual Go > Single Motor; Dual Go > Single Cognitive, respectively, Fig.4). These functionally defined regions of interest (ROIs) then served as the basis for the subsequent task-based functional connectivity analysis (“ROI-to-ROI functional connectivity analysis”). In this analysis, we identified differences in task-based functional connectivity between younger and older adults (Fig. 5 and Fig. 6). In the third step, we examined how individual differences in connectivity related to dual-task behavioral performance using regularized generalized canonical correlation analysis (“Multivariate brain– behavior association analysis”, Fig. 7 and Fig. 8).

#### 2.6.1 Task based GLM analysis and ROI selection

Functional and anatomical MRI data were preprocessed using *fMRIPrep* (Esteban et al., 2019) (version 24.1.1; RRID:SCR_016216), a fully reproducible and standardized preprocessing pipeline designed for task-based and resting-state fMRI analyses (https://fmriprep.org/en/stable/). The preprocessing workflow included fieldmap-based susceptibility distortion correction, head motion correction (realignment), coregistration of functional and anatomical images, brain extraction, tissue segmentation, and spatial normalization to MNI152NLin2009cAsym space. Full details of the preprocessing procedures are provided in the Supplemental Material (Supplementary Material A). Following preprocessing with *fMRIPrep*, the functional data were spatially smoothed using SPM12 (Wellcome Centre for Human Neuroimaging, UCL, London, UK. Available at: https://www.fil.ion.ucl.ac.uk/spm/, (Friston et al., 2007)) with an 8 mm full-width at half-maximum (FWHM) Gaussian kernel to enhance the signal-to-noise ratio for task-based analyses.

At the first-level analysis, five task regressors were specified to model trial-related BOLD responses across cognitive, motor, and dual-task conditions. These included: (i) Single cognitive Go (Go trials in the single-task cognitive); (ii) Single cognitive NoGo (NoGo trials in the single-task cognitive); (iii) Single motor (stepping trials in the single-task motor); (iv) Dual Go (dual-task: Go trials with concurrent stepping); (v) Dual NoGo (dual-task: NoGo with concurrent stepping). Each trial was modeled as a brief event and convolved with the canonical hemodynamic response function (HRF) to estimate condition-specific BOLD responses. Statistical analysis was performed within the general linear model framework (Friston et al., 1994). Six realignment-derived head motion parameters (three translations, three rotations) were included as nuisance regressors to account for residual motion artefacts. To reduce physiological and scanner-related noise, a high-pass filter with a 1/128 Hz cutoff was applied, and a first-order autoregressive (AR(1)) model was applied to correct for temporal autocorrelations (Friston et al., 1995).

To define task-relevant brain networks for subsequent functional connectivity analyses, we computed two targeted contrasts. The dual-task cognitive component contrast (Dual Go > Single Motor) isolated the demands of the cognitive task during dual-tasking, including dual-task control processes. Conversely, the dual-task motor component contrast (Dual Go > Single Cognitive Go) captured the demands of the motor task during dual-tasking including dual-task control processes (see Fig. 4). These contrasts were used solely to identify potential brain regions associated with the cognitive and motor components in the context of dual tasking. Whole-brain activation maps for both single-and dual-task contrasts are reported in the Supplement Fig.S1.

Contrast images were entered into a random-effects general linear model (GLM) on second level to assess task-related activation patterns across all participants (young and older adults combined). Statistical parametric t-maps were generated, and multiple comparisons were controlled using a family-wise error (FWE) correction across the whole brain. A significance threshold of *p* < 0.05 (FWE-corrected) with a minimum cluster extent of 20 contiguous voxels was applied to identify regions with statistically significant activation. Significant clusters for the two targeted contrasts were then extracted as ROIs for subsequent task-based functional connectivity analysis.

To improve anatomical precision, each cluster was spatially constrained using the Harvard-Oxford Cortical and Subcortical Structural Atlas: only voxels within the atlas-defined anatomical boundaries were retained. This means that clusters identified from statistical contrasts were refined by limiting them to voxels strictly within well-defined anatomical regions, ensuring that the resulting ROIs more accurately correspond to known brain structures. Subclusters smaller than 20 voxels after atlas masking were excluded from further analysis. This procedure resulted in 47 ROIs from the dual-task cognitive component contrast and 70 ROIs from the dual-task motor component contrast, which were subsequently entered into ROI-to-ROI functional connectivity analysis in the CONN toolbox(Alfonso Nieto-Castanon, 2020; Whitfield-Gabrieli & Nieto-Castanon, 2012). The full list of ROIs, including anatomical labels and voxel counts, is provided in Supplemental Tab. S1.

#### 2.6.2 Functional Connectivity Analysis

##### Preprocessing

Functional data preprocessing was conducted using the CONN toolbox (Alfonso Nieto-Castanon, 2020; Whitfield-Gabrieli & Nieto-Castanon, 2012) (RRID:SCR_009550, version 22.v2407) and SPM12 (RRID:SCR_007037, release 12.7771) within MATLAB 2023a, performed on a high-performance computing (HPC) cluster. The default preprocessing pipeline in CONN was applied, which includes the following steps: functional realignment and unwarping (with correction for susceptibility distortion interactions), slice-timing correction (to middle slice), outlier detection, segmentation and normalization to standard space, and spatial smoothing. Tissue segmentation was performed directly on each participant’s structural T1-weighted image using the unified segmentation approach in SPM12, which simultaneously segments the image into gray matter, white matter, and CSF compartments while computing nonlinear spatial normalization parameters. These parameters were then applied to normalize both structural and functional images. All images were normalized to MNI152 standard space (Montreal Neurological Institute), specifically the MNI152 nonlinear asymmetrical template (MNI152NLin6Asym) at 2 mm isotropic resolution. Movement outliers were identified using the Artefact Detection Tools (ART) with default thresholds: framewise displacement exceeding 0.9 mm and global signal change exceeding 5 standard deviations. Spatial smoothing was applied using a Gaussian kernel with 8 mm full-width at half-maximum (FWHM).

##### Denoising

Functional data were denoised using the standard denoising pipeline implemented in the CONN toolbox, which involves three key sequential steps: (1) identification of noise-related signal components, (2) linear regression to remove their influence, and (3) temporal band-pass filtering. The denoising model included the following nuisance regressors: five aCompCor components each from white matter and CSF time series (total 10 physiological noise regressors), 12 motion-related parameters (six head motion estimates and their first-order temporal derivatives), and task-related effects modelled as boxcar regressors (convolved with the canonical hemodynamic response function) along with their first-order temporal derivatives (10 regressors in total). Outlier scans were identified using framewise displacement above 0.9 mm or global signal change thresholds (5 STD) via the ART toolbox. Each outlier timepoint (mean: 22 scans per subject out of 1,090 total scans) was entered as a separate nuisance regressor with a value of 1 at the corresponding scan and 0 elsewhere, allowing them to be regressed out independently during the denoising step. Task activation regression was performed to remove condition-related variance from the BOLD signal, ensuring that functional connectivity estimates reflected condition-modulated interactions rather than shared evoked responses. After nuisance regression, a band-pass filter (0.008–0.09 Hz) was applied to isolate low-frequency fluctuations. The effective degrees of freedom in the denoised time series ranged from 94.5 to 147.2 across subjects (mean: 143.1), accounting for the number of regressors and censored scans.

##### ROI-to-ROI Functional Connectivity Analysis

###### First-level analysis

ROI-to-ROI connectivity (RRC) matrices were computed across the 47 and 70 ROIs derived from dual-task cognitive component contrast and the dual-task motor component contrast, respectively, as identified in the section 2.6.1.

Functional connectivity was estimated using a weighted ROI-to-ROI Connectivity (wRRC) measure (Alfonso Nieto-Castanon, 2020), which characterizes task-specific functional connectivity strength (i.e. functional connectivity during each cognitive or motor task, single and dual) among a pre-defined set of ROIs (i.e. cognitive- and motor-ROIs). wRRC matrices are computed using a weighted Least Squares (WLS) linear model with defined temporal weights identifying each individual task condition. In our event-related task design, weights are defined as a task-specific (single- and dual-tasks) boxcar timeseries convolved with a canonical hemodynamic response function. This allows for the estimation of task-specific connectivity, emphasizing periods of active task engagement and reducing the influence of unrelated or baseline fluctuations.

###### Second-level analysis

Group-level analyses were performed using a General Linear Model (Alfonso Nieto-Castanon, 2020). For each individual connection a separate GLM was estimated, with first-level connectivity measures at this connection as dependent variables (one independent sample per subject and one measurement per task condition), and age groups as independent variables. Because each subject contributed multiple connectivity measures across the five task conditions *(Single Cognitive Go, Single Cognitive NoGo, Single Motor Stepping, Dual Go, Dual NoGo)*, group-level analyses accounted for the non-independence of these repeated measures by estimating the covariance structure across these five conditions. Connection-level hypotheses were then evaluated using multivariate parametric statistics with random-effects across subjects and correction for within-subject covariance when these five multiple measurements were present. Statistical inferences were conducted at the cluster level, where clusters represented groups of similar ROI-to-ROI connections. Cluster-level effects were evaluated using parametric statistics both within and between pairs of functional networks (Functional Network Connectivity, (Jafri et al., 2008)). Networks were defined using a complete-linkage hierarchical clustering procedure, which grouped ROIs according to a combination of anatomical proximity and functional similarity metrics (Alfonso Nieto-Castanon, 2020). To control for multiple comparisons, results were thresholded using a two-step approach: first applying a connection-level threshold of *p* < 0.05 (uncorrected), and then applying a cluster-level false discovery rate correction (*p*-FDR < 0.05) to control for multiple comparisons.

To localize age-related effects within brain regions associated with cognitive and motor components of dual-tasking, we conducted domain-specific functional connectivity analyses using two ROI sets defined in the activation’s analysis (see above: Dual Go > Single Motor, and Dual Go > Single Go). Functional connectivity was assessed for each ROI set under both single-and dual-task conditions.

### 2.7 Multivariate brain–behavior association analysis

The aim of this final step was to investigate multivariate associations between functional connectivity and behavioral performance under dual-task conditions, thereby providing additional insights for interpretation. To identify these associations while mitigating multicollinearity and overfitting, we applied regularized Generalized Canonical Correlation Analysis (RGCCA) using the mixOmics R package (v6.22.0; Rohart et al., 2017). RGCCA is a multiblock extension of classical canonical correlation analysis (CCA) that integrates multiple datasets (blocks) measured on the same set of individuals. The method extracts latent variables (components) from each block that maximize correlations across blocks, while incorporating regularization to enhance stability in high-dimensional settings. Subject-level component scores were obtained from this RGCCA analysis, separated by task network and age group. Components represent latent dimensions that maximize shared variance across functional connectivity and behavioral blocks, allowing visualization of group differences in multivariate brain–behavior patterns.

To capture network-specific effects, RGCCA analyses were conducted separately for the **dual-task cognitive component network** and **dual-task motor component network**. Each analysis integrated three data blocks: (1) dual-task functional connectivity derived from the wRRC functional connectivity analysis, (2) single-task (cognitive and motor) functional connectivity derived from the wRRC functional connectivity analysis, and (3) behavioral performance metrics during the dual-task, including RT, variability, error rate and dual-task costs for both motor and cognitive modalities. In the RGCCA block design, the two brain blocks were fully connected to each other and to the behavioral block, enabling examination of multivariate associations both between functional connectivity and behavior, and between single- and dual-task connectivity.

In RGCCA, the regularization parameter tau controls the shrinkage of covariance matrices for each block. Rather than relying on cross-validation, the RGCCA package uses the automatic shrinkage estimator of Schäfer & Strimmer (2005), which analytically derives an optimal shrinkage intensity that balances bias and variance. This yields consistent, well-conditioned covariance matrices without the computational burden of cross-validation. Following Tenenhaus et al. (2017), this method (tau = “optimal”) is the default in RGCCA with τ estimated directly from sample data for each block. The approach is particularly advantageous when the number of variables exceeds the number of subjects, as in our study, where cross-validation can be unstable in small-sample behavioral datasets. Importantly, tau in RGCCA is not tuned for predictive accuracy but to stabilize covariance estimation for multiblock component correlation analysis. Thus, the Schäfer–Strimmer estimator provides a principled, reproducible, and efficient alternative to cross-validation.

## 3. Results

### 3.1 Behavioral results

Older and younger adults did not differ significantly in sex distribution or years of education. As assessed by the TAP-M test battery, older adults performed significantly slower than younger adults in the distraction task, divided attention task (both auditory and visual modalities), and the Go/NoGo task. No significant group difference was observed in the executive control task. To control for multiple comparisons across the five cognitive variables, Bonferroni-adjusted *p*-values were calculated. Descriptive statistics and test results are shown in Tab. 1.

**Table 1.** Participant Characteristics and TAP-M Performance.

| Measure | Older adults<br>(n = 40) | Younger adults<br>(n = 20) | $p$-value |
| --- | --- | --- | --- |
| Age (years) | 67.55 ± 7.09 | 28.00 ± 4.87 | — |
| Gender (male %) | 19 (47.5%) | 11 (55.0%) | 0.784 |
| Education (years) | 13.2 ± 3.09 | 14.6 ± 1.53 | 0.161 |
| **TAP-M** |  |  |  |
| Distraction (ms) | 490.7 ± 9.78 | 414.8 ± 11.34 | **0.0004** |
| Executive Control (ms) | 662.2 ± 16.99 | 623.1 ± 13.83 | 0.396 |
| Divided Attention-auditory (ms) | 651.7 ± 18.60 | 579.4 ± 14.51 | **0.017** |
| Divided Attention-visual (ms) | 854.8 ± 20.12 | 734.2 ± 13.48 | **0.00003** |
| Go/NoGo (ms) | 448.2 ± 10.08 | 388.2 ± 29.17 | **0.0003** |

We first examined RTs to assess how dual-task demands influenced motor and cognitive performance in younger and older adults. RTs differed significantly between younger and older adults (main effect age group: *F*(1, 52) = 4.22, p = .045, ηp² = .044), between single and dual-tasks conditions (main effect task: *F*(1, 52) = 275.10, p < .001, ηp² = .303), and between cognitive and motor domain (main effect of domain: *F*(1, 52) = 173.15, p < .001, ηp² = .433). No significant two-way interactions were observed. Importantly, there was a significant three-way interaction between age, task, and domain (*F*(1, 52) = 11.79, p = .001, ηp² = .027), indicating that dual-task demands affected cognitive and motor performance differently between younger and older adults (see Fig.3).

**Figure 3.**
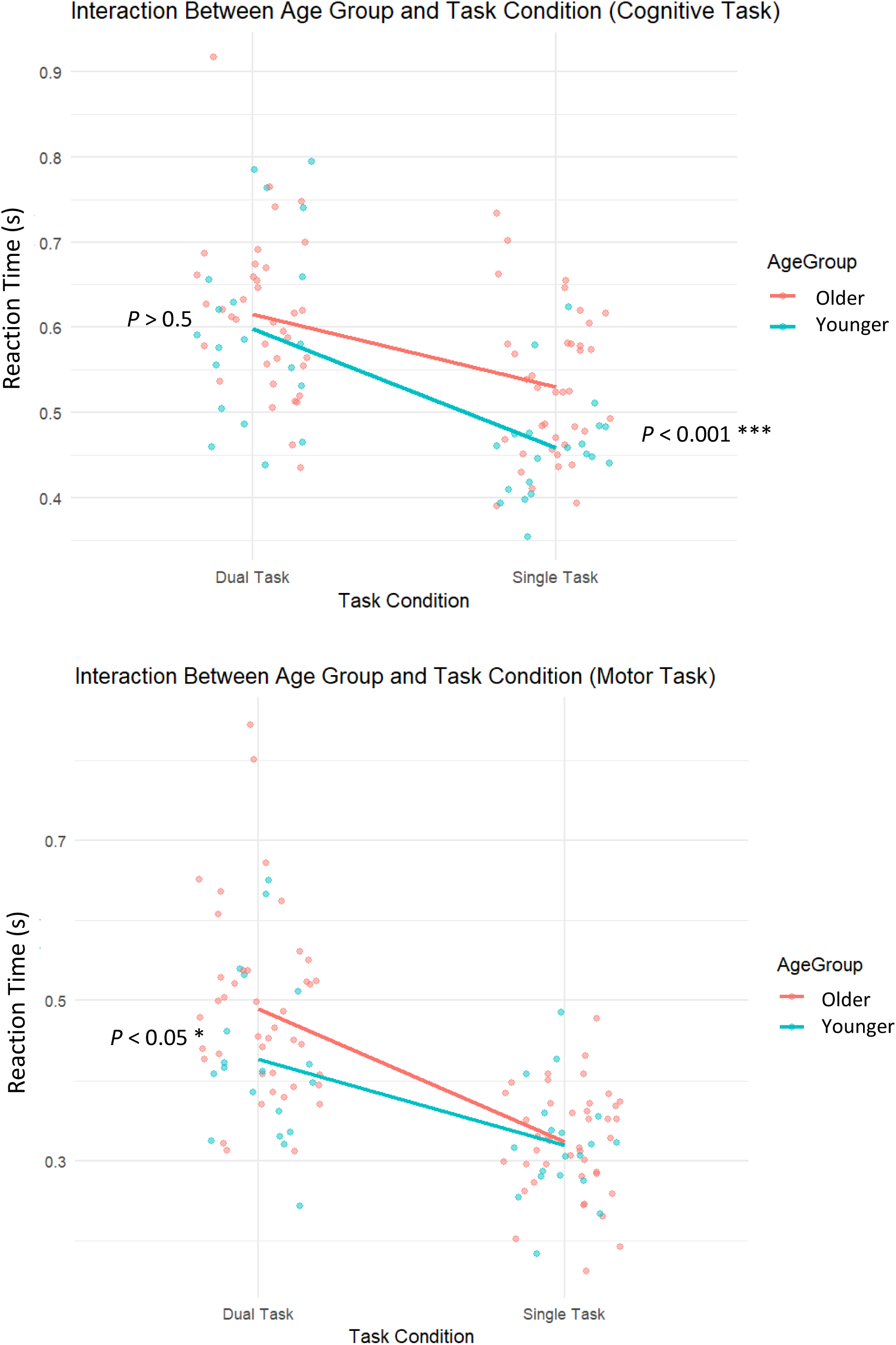
Reaction times (RTs) as a function of age (older vs. younger) and task (dual vs. single) for the cognitive (upper panel) and motor domain (lower panel). In the cognitive domain, older adults (red) responded significantly slower RTs than younger adults (blue) in single-task conditions, but this age difference disappeared under dual-task conditions. In contrast, in the motor domain, older and younger adults performed similar in the single-task condition, but older adults showed significantly longer RTs than younger adults during dual-task performance. Dots represent individual participants, and line plots depicts group means.

As shown in Tab. 2, older adults were slower than younger adults in the cognitive single-task, but this age difference disappeared under dual-task conditions, resulting in comparable cognitive dual-task performance across groups. In contrast, all motor measures revealed pronounced dual-task costs in older adults, including slower responses, greater variability, and higher error rates.

**Table 2.** Mean reaction time (RT), RT variability (SD), and error rate are reported for younger and older adults across single- and dual-task conditions and motor and cognitive domains, including derived dual-task costs (16 outcome measures in total). Two-sample t-tests were used to compare groups for each outcome measure. Values represent mean ± standard error (SE). Bold *p*-values indicate significance at Bonferroni-corrected threshold (*p* < 0.0031).

|  |  | Domain |  |  |  |  |  |
| --- | --- | --- | --- | --- | --- | --- | --- |
| Task | Measure | Cognitive task |  |  | Motor task |  |  |
| | | Old | Young | $p$ | Old | Young | $p$ |
| Single | Mean RT (s) | 0.535 ± 0.014 | 0.459 ± 0.014 | **3.1e-4** | 0.324 ± 0.011 | 0.319 ± 0.016 | 0.827 |
| Dual | Mean RT (s) | 0.617 ± 0.016 | 0.599 ± 0.024 | 0.528 | 0.496 ± 0.018 | 0.426 ± 0.025 | 0.030 |
| | Dual task cost (%) | 15.8 ± 2.69 | 31.4 ± 4.96 | 0.009 | 55.3 ± 4.57 | 33.9 ± 4.36 | **0.001** |
| Single | Variability (s) | 0.085 ± 0.004 | 0.083 ± 0.005 | 0.690 | 0.112 ± 0.004 | 0.107 ± 0.006 | 0.539 |
| Dual | Variability (s) | 0.128 ± 0.005 | 0.113 ± 0.006 | 0.075 | 0.168 ± 0.012 | 0.111 ± 0.005 | **5.5e-5** |
| | Dual task cost (%) | 56.2 ± 7.87 | 44.3 ± 13.3 | 0.447 | 55.3 ± 11.6 | 8.4 ± 6.70 | **9.2e-4** |
| Single | Error Rate (%) | 2.58 ± 0.61 | 5.28 ± 1.11 | 0.038 | 0.10 ± 0.05 | 0.00 ± 0.00 | 0.044 |
| Dual | Error Rate (%) | 3.83 ± 0.82 | 6.30 ± 1.39 | 0.134 | 4.24 ± 1.25 | 0.22 ± 0.10 | **0.0026** |

For variability, all three main effects were significant: age (*F*(1, 52) = 7.27, p = .009, ηp² = .058), task (F(1, 52) = 37.10, p < .001, ηp² = .153), and domain (*F*(1, 52) = 27.12, p < .001, ηp² = .071). In addition, we observed significant two-way interactions between age and task (*F*(1, 52) = 8.55, p = .005, ηp² = .040) and between age and domain (*F*(1, 52) = 7.79, p = .007, ηp² = .022), whereas the task × domain interaction was not significant. Crucially, the three-way interaction of age group, task condition, and domain was also significant (*F*(1, 52) = 6.59, p = .013, ηp² = .020), indicating that performance variability increased most strongly in older adults during motor dual-tasking, whereas variability in the cognitive domain remained largely comparable across groups (see Tab. 2). A similar pattern was seen for errors.

### 3.2 Cognitive and motor components during dual tasking (ROI definition)

To identify regions supporting the cognitive and motor aspects of dual-task performance for subsequent connectivity analyses, we used two targeted contrasts: the dual-task cognitive component contrast (Dual Go – Motor single) and the dual-task motor component contrast (Dual Go – Cognitive Go). The dual-task cognitive component contrast revealed activation primarily in prefrontal, and parietal areas, and occipital cortices, see Fig. 4A, whereas the dual-task motor component contrast was associated with activity in bilateral precentral and postcentral gyrus, superior parietal lobe, basal ganglia and cerebellum, see Fig. 4B. While these patterns partly overlap, as expected given the integrative nature of dual-tasking, they nevertheless allowed us to define ROI masks and examine connectivity separately in networks more strongly associated with either the cognitive or the motor component.

**Figure 4.**
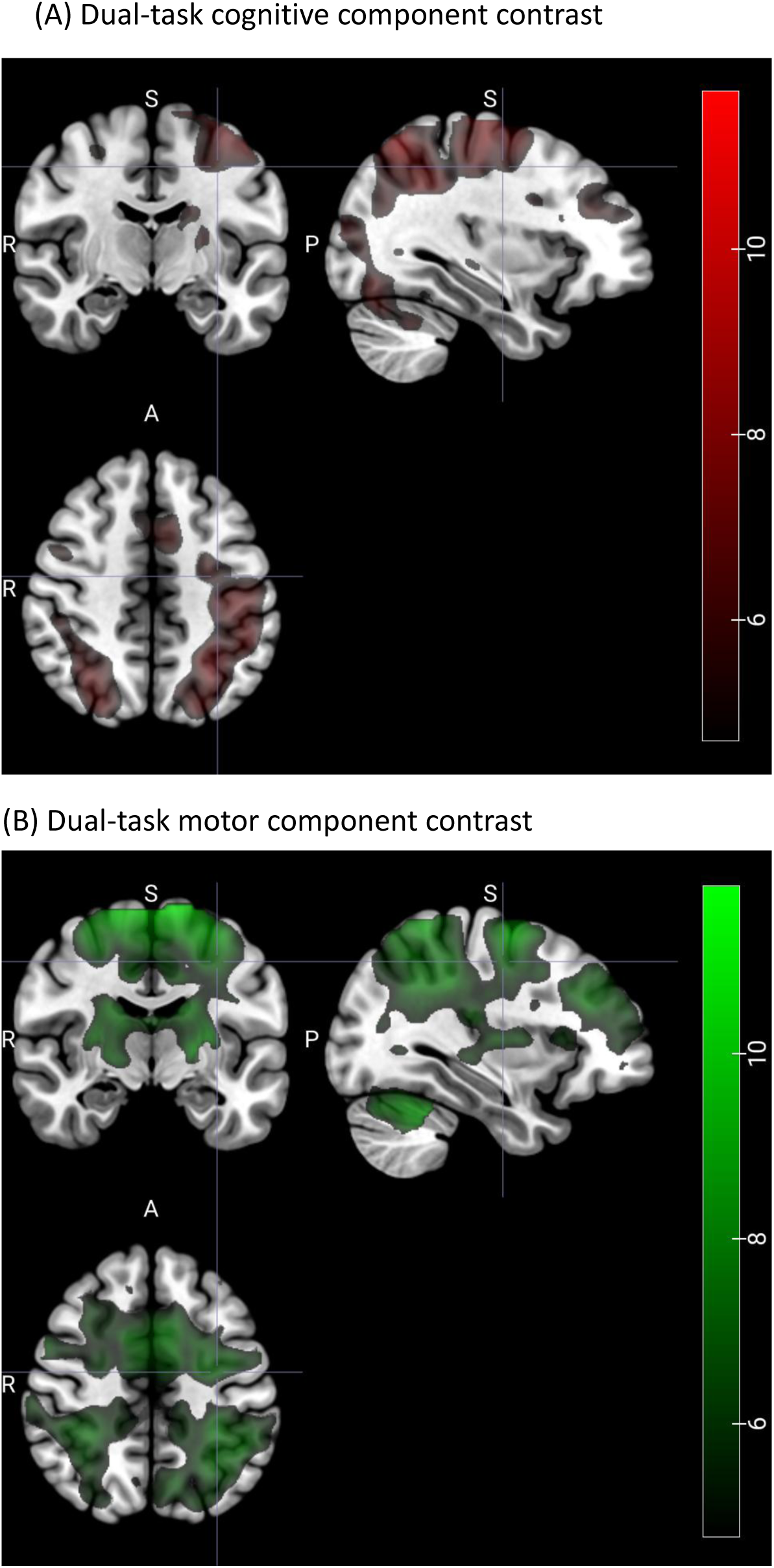
Brain regions identified from contrasts emphasizing the cognitive and motor components of dual-tasking, used for ROI definition in the connectivity analyses. Shown are group-level whole-brain activation patterns for (A) the dual-task cognitive component contrast (Dual Go – Motor single) and (B) the dual-task motor component contrast (Dual Go – Cognitive Go), across age groups, thresholded at p < 0.05 (FWE-corrected).

### 3.3 Age-related differences in functional connectivity within cognitive and motor networks

Building on the behavioral evidence for age-related differences in dual-task costs for motor and cognitive performance, we next examined whether functional connectivity showed corresponding differences within the dual-task cognitive and motor component networks. Since age effects were already evident in the cognitive domain under single-task conditions, connectivity analyses were conducted separately for each task (dual-task and the respective single-task) and domain (cognitive and motor within the ROIs shown in Fig.5 and Fig.6). This approach allowed us to examine whether age-related reorganization of network interactions emerges primarily during dual-task demands or is already present during single-task performance, and whether such effects are more pronounced in cognitive or motor networks.

Within the dual-task cognitive component network, dual-tasking was associated with a limited set of connections that were significantly stronger in older compared to younger adults (Fig. 5A). These involved frontal executive regions such as bilateral superior frontal gyrus (SFG), left frontal pole (FP_l), bilateral middle frontal gyrus (MidFG), as well as motor-planning areas such as the supplementary motor area (SMA) and right precentral gyrus (PreCG_r). Additional involvement of the paracingulate cortex (PaCiG) and left anterior supramarginal gyrus (aSMG_l) was also observed. In contrast, during the single cognitive task, more widespread increases in connectivity in older adults were observed (Fig. 5B), encompassing bilateral frontal and parietal areas including the superior and middle frontal gyri, supramarginal gyrus (SMG), and superior parietal lobules (SPL), as well as several cerebellar clusters.

**Figure 5.**
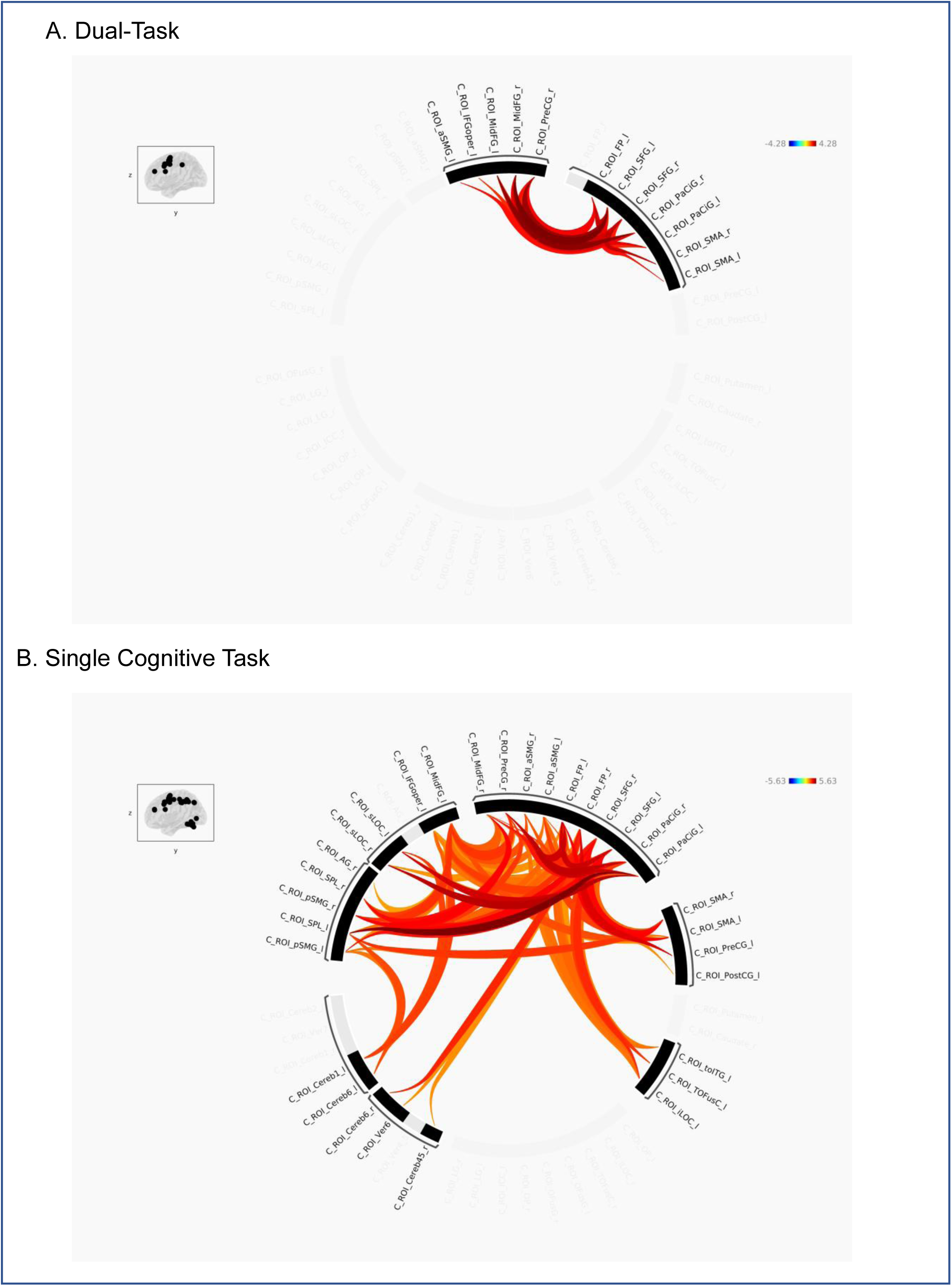
Group-level ROI-to-ROI functional connectivity within the dual-task cognitive component network. Shown are group differences for (A) the dual task, and (B) the single cognitive task. Significant effects (Old > Young; *p* < 0.05, FDR-corrected at the cluster level) are visualized as circular graphs, with color gradients representing t-values (warmer colors: stronger connectivity in older adults; cooler colors: stronger connectivity in younger adults). These results are also presented as a matrix plot in Supplementary Figure S2.

Within the dual-task motor component network, dual tasking was associated with a broader and more distributed pattern of age-related differences in task-related connectivity (Fig. 6A). The areas extended beyond frontal and parietal regions to include posterior cortical areas (e.g., precuneus, occipital cortex), subcortical structures (thalamus, putamen), and the cerebellum. The connectivity profile comprised both positive (Old > Young) and negative (Old < Young) effects. Positive effects were primarily observed within frontal and parietal ROIs, indicating increased connectivity in older adults, whereas negative effects were concentrated in posterior and subcortical regions—particularly between the cerebellum and parietal cortex— indicating reduced coupling between sensory-motor pathways.

In single motor task, age-related effects in functional connectivity were extensive, encompassing both cortical and subcortical regions (Fig. 6B). Older adults exhibited increased functional connectivity across extended bilateral frontal and sensorimotor regions, including the precentral and postcentral gyri, SMA, and basal ganglia. At the same time, reductions in connectivity were observed between occipital, posterior parietal, and cerebellar regions. These findings suggest that even during a relatively simple motor task, aging is associated with widespread network alterations.

**Figure 6.**
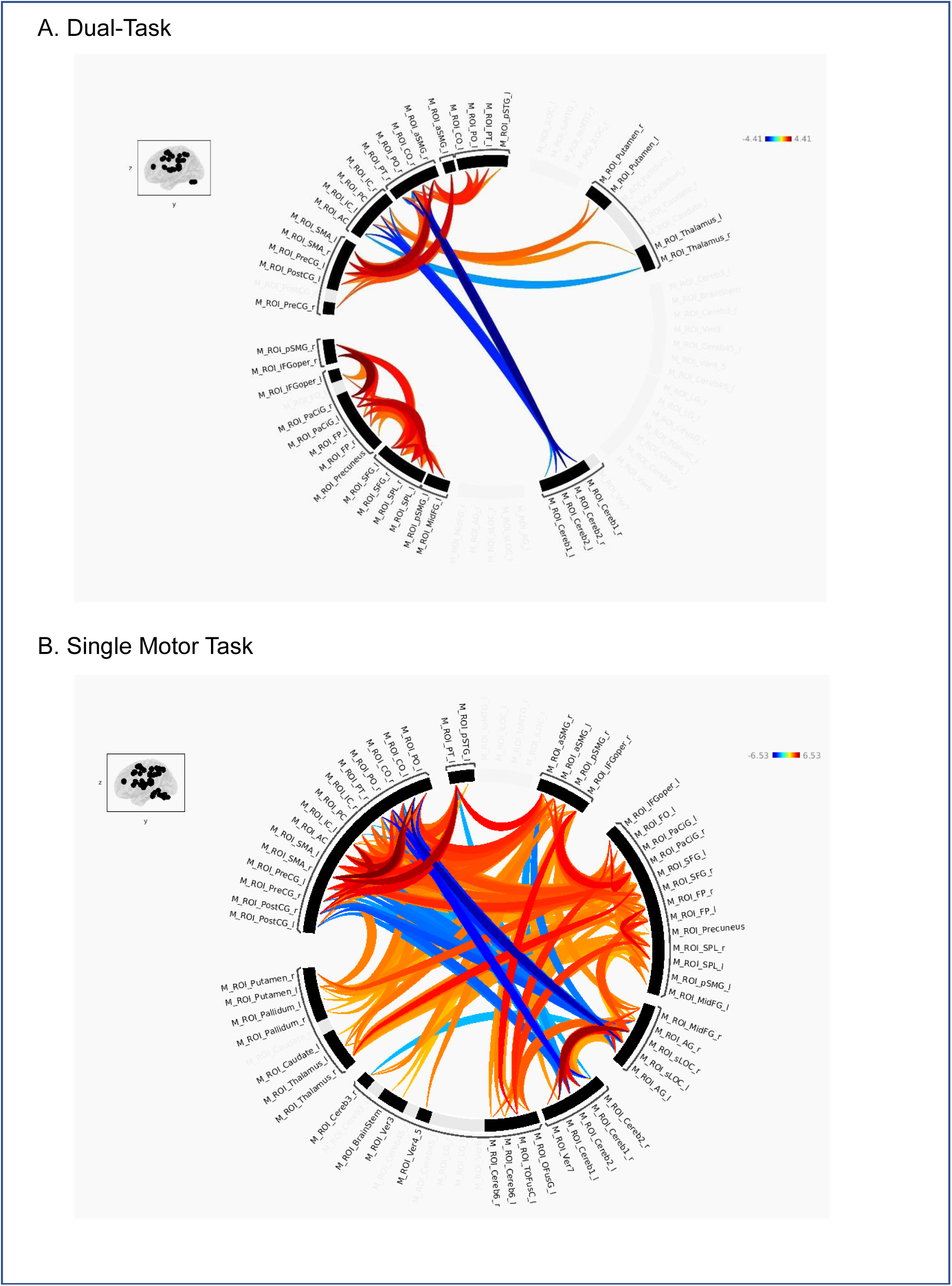
Group-level ROI-to-ROI functional connectivity within the dual-task motor component network. Shown are group differences for (A) the dual-task, and (B) the single motor task. Significant effects (Old > Young; *p* < 0.05, FDR-corrected at the cluster level) are visualized as circular graphs, with color gradients representing t-values (warmer colors: stronger connectivity in older adults; cooler colors: stronger connectivity in younger adults). These results are also presented as a matrix plot in Supplementary Figure S3.

### 3.4 Linking functional connectivity with behavioral performance

To investigate how individual differences in task-based functional connectivity relate to behavioral performance, we conducted multivariate brain–behavior analyses using regularized RGCCA. This approach integrated connectivity measures from both dual- and single-task conditions with a restricted set of behavioral variables collected under dual-task conditions (see Section 2.7). Analyses were performed across the combined sample of younger and older adults, focusing on shared dimensions of covariation between neural and behavioral measures. Fig. 7 shows subject-level component scores derived from the RGCCA analysis, separated by task network and age group. These components represent latent dimensions that maximize shared variance across functional connectivity and behavioral blocks, enabling visualization of group differences in multivariate brain–behavior associations.

The upper panel (Fig. 7A) shows results for the dual-task cognitive component network, including dual-task functional connectivity, single-task functional connectivity (cognitive), and dual-task behavioral performance. The lower panel (Fig. 7B) presents corresponding results for the motor component network, incorporating dual-task functional connectivity, single-task functional connectivity (motor), and dual-task behavior. Participant scores are projected onto the first two RGCCA components, revealing distinct spatial patterns across the two networks. In the motor component network (Fig. 7B), subject scores on Component 1 showed a clear tendency for age-related separation: older adults clustered toward the lower end, whereas younger adults were distributed across a broader and higher range. In contrast, the cognitive component network (Fig. 7A) exhibits a more compressed distribution along both components. Although group differences remained visible along Component 1, the separation was less pronounced, with confidence ellipses of the two age groups showing greater overlap, particularly in dual-task functional connectivity. This reflects a reduced multivariate distance between age groups in cognitive-related functional connectivity and behavior.

Since component 1 showed greater variability and clearer age-related separation, we focused on it to examine correlations among the three data blocks. As shown in Fig. 8, both motor and cognitive networks exhibited significant associations between dual-task performance and functional connectivity under both dual-task and single-task conditions. Compared with younger adults, older adults showed slightly stronger correlations between single-and dual-task functional connectivity and dual-task performance in both networks, except for the single motor functional connectivity and dual-task performance coupling. Notably, FC during the single-task condition correlated with behavior as strongly as during the dual-task condition (e.g., Motor: *r* = .486 vs. .432; Cognitive: *r* = .465 vs. .402), suggesting that connectivity during simpler tasks may capture functional dynamics beyond baseline activity. Diagonal density plots further indicated partial overlap between older and younger adults in the DT cognitive component network, whereas the dual-task motor component network showed a clearer age-related separation.

**Figure 7.**
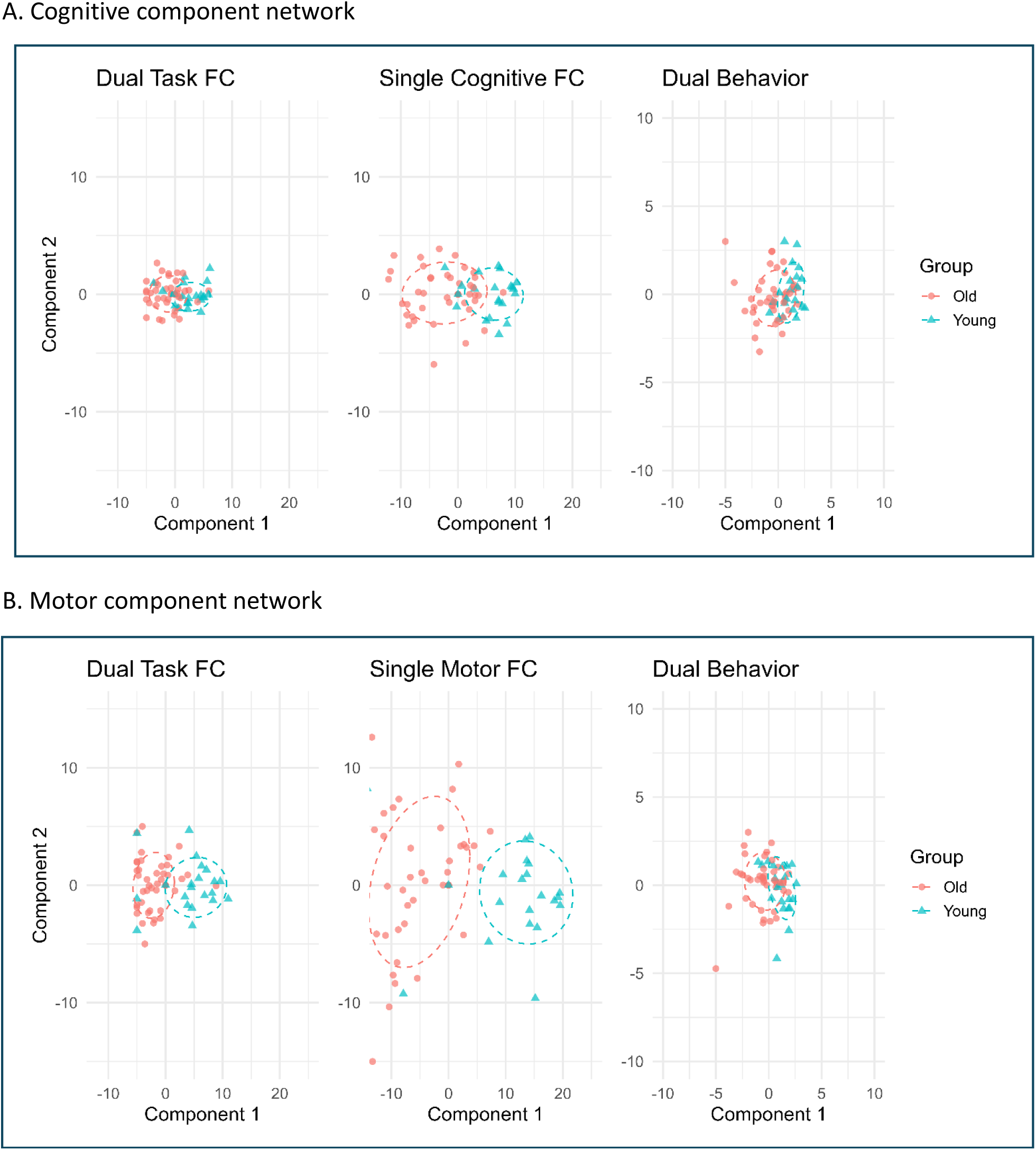
Multivariate component scores from RGCCA analysis by task network and age. Panel A displays component scores within the cognitive task network (Dual-Task FC, Single Cognitive FC, Dual Behavior), while panel B shows scores within the motor network (Dual-Task FC, Single Motor FC, and Dual Behavior). Each plot displays Component 1 versus Component 2 for younger adults (blue triangles) and older adults (red circles), with 68% confidence ellipses indicating group dispersion. For better visual comparison, this figure is also plotted with the axis limits tailored to the scale of each data block, see Supplement Fig. S4. FC: functional connectivity.

**Figure 8.**
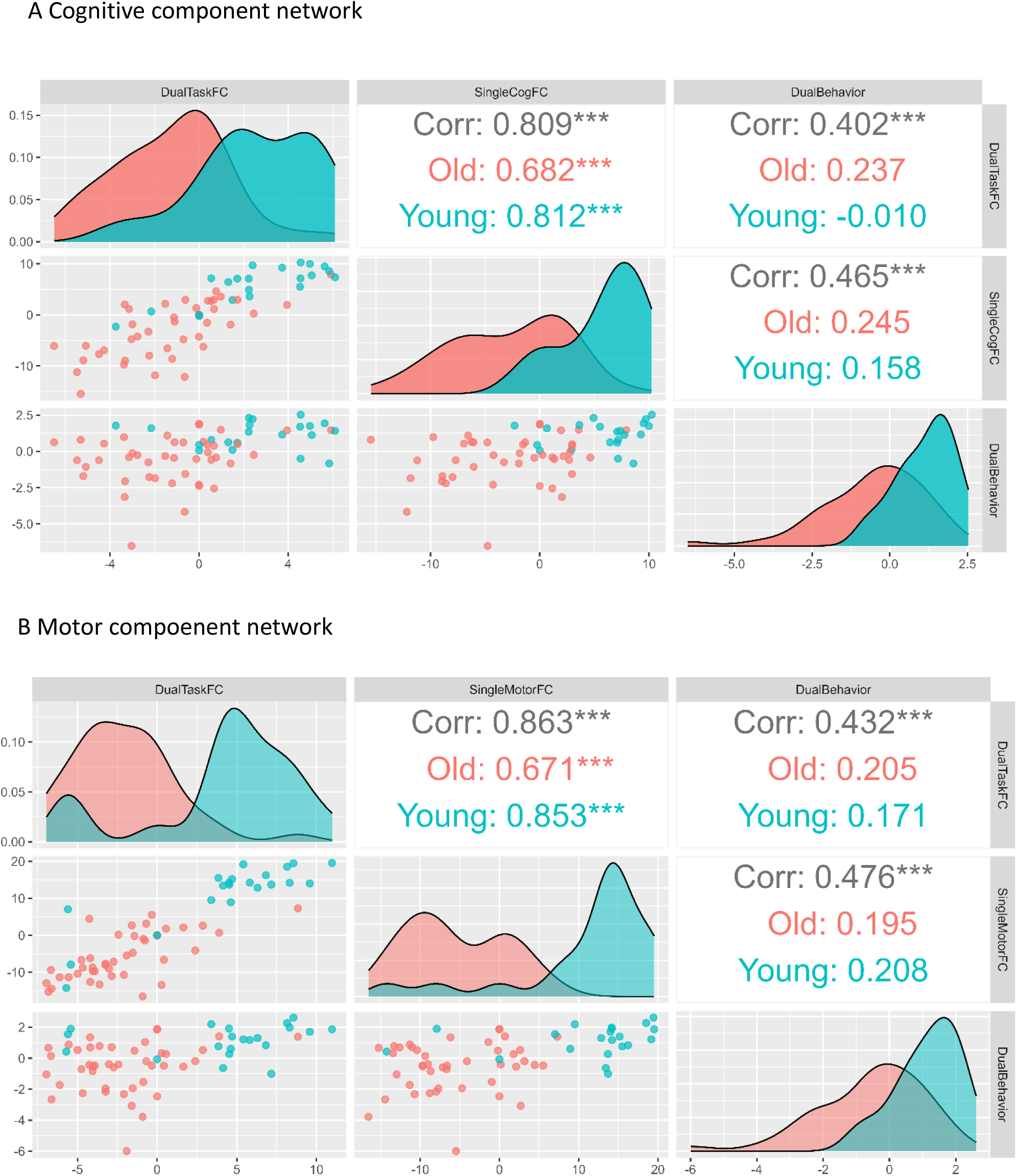
Pairwise correlations for Component 1 scores from the RGCCA analysis of (A) the cognitive component network and (B) the motor component network. Scatterplots (lower panels) and density plots (diagonal) show distributions and relationships for Dual-Task FC, Single-Task FC, and Dual-Task Behaviors. Overall and group-specific correlations (older adults in red, younger adults in blue) are reported in the upper panels. Asterisks indicate significance levels (*** *p* < .001). FC: functional connectivity.

## 4. Discussion

In this study, we investigated task-based functional connectivity during cognitive-motor dual-tasking in older and younger adults using an MRI-compatible pedaling paradigm. Three main findings emerged. First, older adults showed preserved cognitive dual-task performance but greater dual-task costs in motor reaction times and variability. Second, age-related differences in functional connectivity were evident not only under dual-task demands but also during the respective single tasks. These effects were network-specific: connectivity changes in cognitive networks were relatively selective involving frontal executive and motor-planning circuits, whereas motor networks showed broader, system-wide alterations including reduced posterior and subcortical sensorimotor coupling with notable involvement of cerebellar nodes. Third, multivariate RGCCA indicated that age is associated with systematic differences in latent connectivity patterns across dual- and single-task performance, which aligned with a latent behavioral dimension.

### 4.1 Domain-specific dual-task costs in aging

Behaviorally, we observed a significant increase in RT for the single cognitive task in the older group, but no significant differences in RT, its variability, or error rate during the dual task. Comparable cognitive performance across age groups in dual-task settings has also been reported by O’Connell & Basak, (2018) and by Plummer-D’Amato et al., (2012) across two different cognitive tasks. Notably, Plummer-D’Amato et al. (2012) demonstrated that lower education is associated with greater cognitive dual-task costs, while it had no effect on gait. In our study, both age groups had relatively high and comparable years of education, which may partly account for their relatively comparable cognitive performance in the dual task. This lack of group differences may also reflect the relative ease of the Go/NoGo cognitive paradigm employed here, potentially reducing sensitivity to age-related decline. This aligns with previous dual-task findings showing that the complexity of the cognitive component critically modulates the magnitude of age effects (Bianchini et al., 2022; Goh et al., 2021).

By contrast, in the motor domain, older adults showed significantly higher dual-task costs across RT, variability, and error rate, with dual-task variability showing the largest effect. This distinct domain-specific pattern was also observed in both fMRI (Bürki et al., 2017) and ground walking studies (Hollman et al., 2007b; Piche et al., 2023). The observed predominantly motor decrement under dual-task conditions aligns with a recent systematic review and meta-analysis of 39 studies on locomotor-cognitive dual-tasking (Mustafovska et al., 2025), which reported that velocity variability during ground walking showed the largest group difference. According to the classification proposed by Collyer et al., (2022), which categorizes older adults into four groups: dual decline (gait and cognition), gait decline only, cognitive decline only, and non-decliners, our findings suggest that the present sample predominantly exhibits motor-only decline, rather than a dual decline pattern encompassing both cognitive and motor functions.

### 4.2 Functional network level reorganization in cognitive and motor domains in aging

Consistent with behavioral findings, our neuroimaging analyses revealed distinct functional connectivity patterns during single and dual-task processing for younger and older adults. In the dual-task cognitive component network, older adults showed widespread increases in functional connectivity, predominantly involving bilateral frontal and parietal regions, such as the superior and middle frontal gyri, SMG, and SPL. These effects were largely positive, indicating increased synchrony within the cognitive control network in older adults compared to young. The findings are in line with prior task-based connectivity studies showing that aging is associated with reorganization of large-scale networks with additional frontal network recruitment during cognitive engagement (Geerligs et al., 2014; Matthäus et al., 2012; O’Connell & Basak, 2018; Stojan et al., 2023). Notably, Geerligs et al. (2014) demonstrated that such increased recruitment in older adults emerges already during relatively simple tasks. Under dual-task conditions, by contrast increases were markedly more focussed and largely confined to frontal executive (e.g., bilateral SFG, left FP, bilateral MidFG) and motor-planning regions (e.g., SMA, precentral gyrus). This pattern suggests a targeted upregulation of frontally mediated cognitive control resources, consistent with compensatory recruitment to manage increased task demands (Cappell et al., 2010; Droby et al., 2022; Matthäus et al., 2012). Behaviorally, this pattern may underpin the finding that older adults were able to maintain dual-task cognitive performance at a level comparable to younger adults. Although the compensation-related utilization of neural circuits hypothesis (CRUNCH) and the scaffolding theory of aging and cognition (STAC) were originally formulated in terms of regional activation (Cabeza et al., 2018; Park & Reuter-Lorenz, 2009), our findings are consistent with their extension to the connectivity level, where compensation is expressed not only through local over-activation but also through strengthened inter-regional coupling (Grady et al., 2016; Penhale et al., 2024). The observed coupling between executive and motor regions may reflect enhanced integration of cognitive control and motor planning mechanisms during cognitive-motor task performance (Dørum et al., 2017; Heuninckx et al., 2008; O’Connell & Basak, 2018).

By contrast, the dual task motor component network revealed a more distributed, system-wide reorganization under the same dual-task condition. Age-related connectivity differences extended beyond frontal and parietal regions to posterior cortical regions (e.g., precuneus, occipital cortex), subcortical structures (thalamus, putamen), and the cerebellum. Notably, older adults exhibited both strengthened (Old > Young) and weakened (Old < Young) connectivity: stronger connectivity emerged in frontoparietal control regions, while reduced connectivity was observed in posterior and cerebellar–parietal pathways. These results suggest that the motor aspect of dual-tasking in aging engages a more extensive reorganization of functional networks, characterized by strengthened top-down control but reduced bottom-up sensorimotor integration. This pattern parallels the behavioral finding of disproportionate motor dual-task slowing and performance instability in older adults, and is consistent with prior work highlighting age-related shifts toward compensatory control and diminished sensorimotor efficiency (Bernard & Seidler, 2013; Seidler et al., 2010).

Importantly, our data highlight the cerebellum as a key locus of reduced network dynamics in older adults. Once framed as a motor structure, the cerebellum is now recognized as a key node for cognition across domains. This broader role is particularly relevant for aging-related conditions such as Parkinson’s and Alzheimer’s, where posterior/vermal atrophy and altered cerebello-cortical coupling often yield early compensatory up-regulation (Liu et al., 2024; McElroy et al., 2024; B. Wang et al., 2025). In the single-task conditions, older adults showed mostly increased cerebellar connectivity (with some regional decreases in the motor task), suggesting that under moderate demands the cerebellum can still provide compensatory support for motor–cognitive integration. This pattern aligns with meta-analytic topography that places sequencing, temporal prediction, and internal-model processing in anterior cerebellar subdivisions and with a modular organization of cerebellar contributions across task domains (Bernard & Seidler, 2013). However, under dual-task conditions—where integrative demands further increase, this pattern reversed: older adults exhibited reduced coupling between the cerebellum and parietal regions and with anterior insular and cingulate cortices. Thus, in older adults the cerebellum’s supportive role may become insufficient when multiple domains must be coordinated simultaneously. This interpretation is consistent with lifespan accounts positing a shift in the cerebellum’s role from crucial in childhood to supportive in adulthood, leaving older adults vulnerable when task demands exceed compensatory reserves (Beuriat et al., 2022). Structurally, a recent diffusion tensor imaging study shows accelerated age-related decline in the cerebello-thalamo-cortical tract, with reduced white-matter integrity predicting poorer executive function from late middle age onward (Kraft et al., 2025). Together with evidence for declining long-range functional hubs and compensatory increases in cerebellar connectivity with age (Tomasi & Volkow, 2012), our findings suggest that the cerebellum is both a site of early adaptive reorganization and a critical bottleneck for maintaining dual-task performance in later life. Our data therefore extend compensatory-recruitment models by highlighting task-dependent vulnerability: supportive contributions are evident under single-task demands, but integrative dynamics degrade under dual-task load.

### 4.3 Latent brain–behavior dimensions reveal age-specific patterns

To examine how reorganization of functional connectivity relates to behavior in an integrated way across multiple measures in our sample, we applied a multivariate approach (RGCCA) (Rohart et al., 2017; Tenenhaus et al., 2017; Welham et al., 2023). This analysis identifies latent components that show maximal correlations between blocks of variables. Interestingly, the first RGCCA component (Fig. 7) captured a shared gradient, along which age groups showed a visible tendency toward separation in a domain- and task-dependent manner. In the dual-task motor component network, older adults displayed greater dispersion in component scores during single-task performance, suggesting heterogeneous reorganization strategies during basic motor execution. By contrast, under dual-task conditions, older adults clustered more tightly at the lower end of the gradient, consistent with more selective and modular reorganization (Cole et al., 2013; Greene et al., 2020). In the dual-task cognitive component network, group differences along Component 1 were less pronounced, with both age groups showing compressed and overlapping score distributions, suggesting more subtle or variable age-related changes in connectivity– behavior coupling. Overall, these patterns tentatively suggest that motor systems may be more vulnerable to aging, whereas cognitive systems may retain more flexible or resilient coupling mechanisms (Grande et al., 2023; Song et al., 2014; Ward & Frackowiak, 2003).

To describe how functional connectivity and behavior jointly align along the latent dimension of component 1 in old and young, we examined the block-specific component scores and their pairwise correlations. In the motor network, younger adults showed particularly strong correlations between single- and dual-task connectivity, whereas these associations were weaker and more variable in older adults. By contrast, correlations with behavioral performance were generally modest and similar across groups. In the cognitive network, associations were overall weaker and largely overlapping between groups. This pattern suggests that age-related separation in the latent space is primarily driven by motor network reorganization, where younger adults show more consistent coupling between FC measures, whereas older adults display greater dispersion and less stable FC–behavior associations. Notably, correlations with single-task FC were comparable in strength to those with dual-task FC, indicating that even in simpler contexts, connectivity captures functional dynamics relevant for dual-task performance. Recent work on functional connectome gradients further underscores that aging affects large-scale network hierarchies in a non-linear, system-specific manner (Riccardi et al., 2025; Tomasi & Volkow, 2012; X. Wang et al., 2024). Our finding that motor networks show greater dispersion and vulnerability, whereas cognitive networks remain relatively preserved, is consistent with these broader observations of gradient dedifferentiation and selective resilience across systems.

## 5. Conclusion

By examining network-level connectivity during cognitive–motor dual-tasking, we provide evidence that cognitive and motor components of those networks show age-related changes in functional connectivity that may reflect resilience and vulnerability. We observed increased variability of functional connectivity in motor networks already during single-task performance in the older group, suggesting that heterogeneity in basic motor-network stability may be a critical factor that determines why some older adults maintain function while others decline. While these findings point to motor-network variability as a promising target for understanding preserved versus declining function in aging and neurodegenerative disease, replication in independent samples and longitudinal designs will be essential to establish its robustness and predictive value.

## 6. Data and code availability

Upon acceptance, all data and analysis code will be uploaded to the Open Science Framework (OSF) and made publicly available.

## 7. Author Contributions

Yan Deng: Methodology, Software, Formal analysis, Visualization, Investigation, Writing – Original draft preparation

AmirHussein Abdolalizadeh: Supervision, Software, Writing – Reviewing and Editing

Kayson Fakhar: Methodology, Investigation, Visualization, Writing – Reviewing and Editing

Carsten Gießing: Supervision, Writing – Reviewing and Editing

Tina Schmitt: Methodology, Writing – Reviewing and Editing

Jochem W. Rieger: Methodology, Writing – Reviewing and Editing

Karsten Witt: Supervision, Writing – Reviewing and Editing

Christiane M. Thiel: Conceptualization, Funding acquisition, Supervision, Writing – Original draft preparation

## 10. Acknowledgements

The multiband EPI sequence (CMRR MB-EPI) was kindly provided by the Center for Magnetic Resonance Research (CMRR), University of Minnesota. We thank Gülsen Yanc and Britta Bruns for their support during MRI scanning.

## References

Alfonso Nieto-Castanon. (2020). Handbook of functional connectivity Magnetic Resonance Imaging methods in CONN. ResearchGate.

Ali, P., Renaud, P., Montero-Odasso, M., Gautier, J., Dinomais, M., & Annweiler, C. (2024). Gait performance in older adults across the cognitive spectrum: Results from the GAIT cohort. Journal of the American Geriatrics Society, 72(11), 3437–3447. 10.1111/jgs.19162

Belghali, M., Chastan, N., Cignetti, F., Davenne, D., & Decker, L. M. (2017). Loss of gait control assessed by cognitive-motor dual-tasks: Pros and cons in detecting people at risk of developing Alzheimer’s and Parkinson’s diseases. GeroScience, 39(3), 305–329. 10.1007/s11357-017-9977-7

Bernard, J. A., & Seidler, R. D. (2013). Cerebellar contributions to visuomotor adaptation and motor sequence learning: An ALE meta-analysis. Frontiers in Human Neuroscience, 7, 27. 10.3389/fnhum.2013.00027

Beuriat, P.-A., Cristofori, I., Gordon, B., & Grafman, J. (2022). The shifting role of the cerebellum in executive, emotional and social processing across the lifespan. Behavioral and Brain Functions, 18(1), 6. 10.1186/s12993-022-00193-5

Bianchini, E., Warmerdam, E., Romijnders, R., Hansen, C., Pontieri, F. E., & Maetzler, W. (2022). Cognitive dual-task cost depends on the complexity of the cognitive task, but not on age and disease. Frontiers in Neurology, 13. 10.3389/fneur.2022.964207

Bishnoi, A., & Hernandez, M. E. (2021). Dual task walking costs in older adults with mild cognitive impairment: A systematic review and meta-analysis. Aging & Mental Health, 25(9), 1618– 1629. 10.1080/13607863.2020.1802576

Bohle, H., Rimpel, J., Schauenburg, G., Gebel, A., Stelzel, C., Heinzel, S., Rapp, M., & Granacher, U. (2019). Behavioral and Neural Correlates of Cognitive-Motor Interference during Multitasking in Young and Old Adults. Neural Plasticity, 2019, 1–19. 10.1155/2019/9478656

Bookheimer, S. Y., Salat, D. H., Terpstra, M., Ances, B. M., Barch, D. M., Buckner, R. L., Burgess, G. C., Curtiss, S. W., Diaz-Santos, M., Elam, J. S., Fischl, B., Greve, D. N., Hagy, H. A., Harms, M. P., Hatch, O. M., Hedden, T., Hodge, C., Japardi, K. C., Kuhn, T. P., … Yacoub, E. (2019). The Lifespan Human Connectome Project in Aging: An overview. NeuroImage, 185, 335–348. 10.1016/j.neuroimage.2018.10.009

Boyne, P., Maloney, T., DiFrancesco, M., Fox, M. D., Awosika, O., Aggarwal, P., Woeste, J., Jaroch, L., Braswell, D., & Vannest, J. (2018). Resting-state functional connectivity of subcortical locomotor centers explains variance in walking capacity. Human Brain Mapping, 39(12), 4831–4843. 10.1002/hbm.24326

Bürki, C. N., Bridenbaugh, S. A., Reinhardt, J., Stippich, C., Kressig, R. W., & Blatow, M. (2017). Imaging gait analysis: An FMRI dual task study. Brain and Behavior, 7(8), e00724. 10.1002/brb3.724

Cabeza, R., Albert, M., Belleville, S., Craik, F. I. M., Duarte, A., Grady, C. L., Lindenberger, U., Nyberg, L., Park, D. C., Reuter-Lorenz, P. A., Rugg, M. D., Steffener, J., & Rajah, M. N. (2018). Maintenance, reserve and compensation: The cognitive neuroscience of healthy ageing. Nature Reviews Neuroscience, 19(11), 701–710. 10.1038/s41583-018-0068-2

Cappell, K. A., Gmeindl, L., & Reuter-Lorenz, P. A. (2010). Age differences in prefontal recruitment during verbal working memory maintenance depend on memory load. Cortex; a Journal Devoted to the Study of the Nervous System and Behavior, 46(4), 462–473. 10.1016/j.cortex.2009.11.009

Cole, M. W., Reynolds, J. R., Power, J. D., Repovs, G., Anticevic, A., & Braver, T. S. (2013). Multi-task connectivity reveals flexible hubs for adaptive task control. Nature Neuroscience, 16(9), 1348–1355. 10.1038/nn.3470

Collyer, T. A., Murray, A. M., Woods, R. L., Storey, E., Chong, T. T.-J., Ryan, J., Orchard, S. G., Brodtmann, A., Srikanth, V. K., Shah, R. C., & Callisaya, M. L. (2022). Association of Dual Decline in Cognition and Gait Speed With Risk of Dementia in Older Adults. JAMA Network Open, 5(5), e2214647. 10.1001/jamanetworkopen.2022.14647

Deary, I. J., Corley, J., Gow, A. J., Harris, S. E., Houlihan, L. M., Marioni, R. E., Penke, L., Rafnsson, S. B., & Starr, J. M. (2009). Age-associated cognitive decline. British Medical Bulletin, 92, 135–152. 10.1093/bmb/ldp033

Dørum, E. S., Kaufmann, T., Alnæs, D., Andreassen, O. A., Richard, G., Kolskår, K. K., Nordvik, J. E., & Westlye, L. T. (2017). Increased sensitivity to age-related differences in brain functional connectivity during continuous multiple object tracking compared to resting-state. NeuroImage, 148, 364–372. 10.1016/j.neuroimage.2017.01.048

Droby, A., Varangis, E., Habeck, C., Hausdorff, J. M., Stern, Y., Mirelman, A., & Maidan, I. (2022). Effects of aging on cognitive and brain inter-network integration patterns underlying usual and dual-task gait performance. Frontiers in Aging Neuroscience, 14, 956744. 10.3389/fnagi.2022.956744

Esteban, O., Markiewicz, C. J., Blair, R. W., Moodie, C. A., Isik, A. I., Erramuzpe, A., Kent, J. D., Goncalves, M., DuPre, E., Snyder, M., Oya, H., Ghosh, S. S., Wright, J., Durnez, J., Poldrack, R. A., & Gorgolewski, K. J. (2019). fMRIPrep: A robust preprocessing pipeline for functional MRI. Nat Methods, 16(1), 111–116. 10.1038/s41592-018-0235-4

Feinberg, D. A., Moeller, S., Smith, S. M., Auerbach, E., Ramanna, S., Gunther, M., Glasser, M. F., Miller, K. L., Ugurbil, K., & Yacoub, E. (2010). Multiplexed echo planar imaging for sub-second whole brain FMRI and fast diffusion imaging. PloS One, 5(12), e15710. 10.1371/journal.pone.0015710

Friston, K. J., Ashburner, J., Kiebel, S., Nichols, T., & Penny, W. (2007). Statistical Parametric Mapping: The Analysis of Functional Brain Images. Academic Press. 10.1016/B978-012372560-8/50003-6

Friston, K. J., Holmes, A. P., Poline, J.-B., Grasby, P. J., Williams, S. C. R., Frackowiak, R. S. J., & Turner, R. (1995). Analysis of fMRI Time-Series Revisited. NeuroImage, 2(1), 45–53. 10.1006/nimg.1995.1007

Friston, K. J., Holmes, A. P., Worsley, K. J., Poline, J.-P., Frith, C. D., & Frackowiak, R. S. J. (1994). Statistical parametric maps in functional imaging: A general linear approach. Human Brain Mapping, 2(4), 189–210. 10.1002/hbm.460020402

Geerligs, L., Saliasi, E., Renken, R. J., Maurits, N. M., & Lorist, M. M. (2014). Flexible connectivity in the aging brain revealed by task modulations. Human Brain Mapping, 35(8), 3788–3804. 10.1002/hbm.22437

Goh, H.-T., Pearce, M., & Vas, A. (2021). Task matters: An investigation on the effect of different secondary tasks on dual-task gait in older adults. BMC Geriatrics, 21(1), 510. 10.1186/s12877-021-02464-8

Grady, C., Sarraf, S., Saverino, C., & Campbell, K. (2016). Age differences in the functional interactions among the default, frontoparietal control, and dorsal attention networks. Neurobiology of Aging, 41, 159–172. 10.1016/j.neurobiolaging.2016.02.020

Grande, G., Vetrano, D. L., Kalpouzos, G., Welmer, A.-K., Laukka, E. J., Marseglia, A., Fratiglioni, L., & Rizzuto, D. (2023). Brain Changes and Fast Cognitive and Motor Decline in Older Adults. The Journals of Gerontology: Series A, 78(2), 326–332. 10.1093/gerona/glac177

Greene, A. S., Gao, S., Noble, S., Scheinost, D., & Constable, R. T. (2020). How Tasks Change Whole-Brain Functional Organization to Reveal Brain-Phenotype Relationships. Cell Reports, 32(8), 108066. 10.1016/j.celrep.2020.108066

Heuninckx, S., Wenderoth, N., & Swinnen, S. P. (2008). Systems neuroplasticity in the aging brain: Recruiting additional neural resources for successful motor performance in elderly persons. The Journal of Neuroscience: The Official Journal of the Society for Neuroscience, 28(1), 91–99. 10.1523/JNEUROSCI.3300-07.2008

Hollman, J. H., Kovash, F. M., Kubik, J. J., & Linbo, R. A. (2007a). Age-related differences in spatiotemporal markers of gait stability during dual task walking. Gait & Posture, 26(1), 113–119. 10.1016/j.gaitpost.2006.08.005

Hollman, J. H., Kovash, F. M., Kubik, J. J., & Linbo, R. A. (2007b). Age-related differences in spatiotemporal markers of gait stability during dual task walking. Gait & Posture, 26(1), 113–119. 10.1016/j.gaitpost.2006.08.005

Holtzer, R., Mahoney, J. R., Izzetoglu, M., Izzetoglu, K., Onaral, B., & Verghese, J. (2011). fNIRS Study of Walking and Walking While Talking in Young and Old Individuals. The Journals of Gerontology Series A: Biological Sciences and Medical Sciences, 66A(8), 879–887. 10.1093/gerona/glr068

Jafri, M. J., Pearlson, G. D., Stevens, M., & Calhoun, V. D. (2008). A method for functional network connectivity among spatially independent resting-state components in schizophrenia. NeuroImage, 39(4), 1666–1681. 10.1016/j.neuroimage.2007.11.001

Kim, J., Rider, J. V., Zinselmeier, A., Chiu, Y.-F., Peterson, D., & Longhurst, J. K. (2024). Dual-task gait has prognostic value for cognitive decline in Parkinson’s disease. Journal of Clinical Neuroscience, 126, 101–107. 10.1016/j.jocn.2024.06.006

Kraft, J. N., Ortega, A., Hoagey, D. A., Rodrigue, K. M., & Kennedy, K. M. (2025). Age-related cerebello-thalamo-cortical white matter degradation and executive function performance across the lifespan (p. 2025.08.22.671779). bioRxiv. 10.1101/2025.08.22.671779

Leone, C., Feys, P., Moumdjian, L., D’Amico, E., Zappia, M., & Patti, F. (2017). Cognitive-motor dual-task interference: A systematic review of neural correlates. Neurosci Biobehav Rev, 75, 348–360. 10.1016/j.neubiorev.2017.01.010

Lino, T. B., Scarmagnan, G. S., Sobrinho-Junior, S. A., Tessari, G. M. F., Gonçalves, G. H., Pereira, H. M., & Christofoletti, G. (2023). Impact of Using Smartphone While Walking or Standing: A Study Focused on Age and Cognition. Brain Sciences, 13(7), 987. 10.3390/brainsci13070987

Liu, G., Yang, C., Wang, X., Chen, X., Cai, H., & Le, W. (2024). Cerebellum in neurodegenerative diseases: Advances, challenges, and prospects. iScience, 27(11), 111194. 10.1016/j.isci.2024.111194

Longhurst, J. K., Rider, J. V., Cummings, J. L., John, S. E., Poston, B., & Landers, M. R. (2023). Cognitive-motor dual-task interference in Alzheimer’s disease, Parkinson’s disease, and prodromal neurodegeneration: A scoping review. Gait & Posture, 105, 58–74. 10.1016/j.gaitpost.2023.07.277

Matthäus, F., Schmidt, J.-P., Banerjee, A., Schulze, T. G., Demirakca, T., & Diener, C. (2012). Effects of age on the structure of functional connectivity networks during episodic and working memory demand. Brain Connectivity, 2(3), 113–124. 10.1089/brain.2012.0077

McElroy, C. L., Wang, B., Zhang, H., & Jin, K. (2024). Cerebellum and Aging: Update and Challenges. Aging and Disease, 15(6), 2345–2360. 10.14336/AD.2024.0220

Moeller, S., Yacoub, E., Olman, C. A., Auerbach, E., Strupp, J., Harel, N., & Uğurbil, K. (2010). Multiband multislice GE-EPI at 7 tesla, with 16-fold acceleration using partial parallel imaging with application to high spatial and temporal whole-brain fMRI. Magnetic Resonance in Medicine, 63(5), 1144–1153. 10.1002/mrm.22361

Montero-Odasso, M., Speechley, M., Muir-Hunter, S. W., Sarquis-Adamson, Y., Sposato, L. A., Hachinski, V., Borrie, M., Wells, J., Black, A., Sejdić, E., Bherer, L., Chertkow, H., & The Canadian Gait and Cognition Network. (2018). Motor and Cognitive Trajectories Before Dementia: Results from Gait and Brain Study. Journal of the American Geriatrics Society, 66(9), 1676–1683. 10.1111/jgs.15341

Mustafovska, J., Wilson, P. H., Cole, M. H., & McGuckian, T. B. (2025). Locomotor-cognitive dual-tasking is reduced in older adults relative to younger: A systematic review with meta-analysis. Gait & Posture, 120, 177–191. 10.1016/j.gaitpost.2025.04.012

Niederer, D., Engeroff, T., Fleckenstein, J., Vogel, O., & Vogt, L. (2021). The age-related decline in spatiotemporal gait characteristics is moderated by concerns of falling, history of falls & diseases, and sociodemographic-anthropometric characteristics in 60–94 years old adults. European Review of Aging and Physical Activity, 18(1), 19. 10.1186/s11556-021-00275-9

O’Connell, M. A., & Basak, C. (2018). Effects of task complexity and age-differences on task-related functional connectivity of attentional networks. Neuropsychologia, 114, 50–64. 10.1016/j.neuropsychologia.2018.04.013

Papegaaij, S., Hortobágyi, T., Godde, B., Kaan, W. A., Erhard, P., & Voelcker-Rehage, C. (2017). Neural correlates of motor-cognitive dual-tasking in young and old adults. PLOS ONE, 12(12), e0189025. 10.1371/journal.pone.0189025

Park, D. C., & Reuter-Lorenz, P. (2009). The Adaptive Brain: Aging and Neurocognitive Scaffolding. Annual Review of Psychology, 60(1), 173–196. 10.1146/annurev.psych.59.103006.093656

Penhale, S. H., Arif, Y., Schantell, M., Johnson, H. J., Willett, M. P., Okelberry, H. J., Meehan, C. E., Heinrichs-Graham, E., & Wilson, T. W. (2024). Healthy aging alters the oscillatory dynamics and fronto-parietal connectivity serving fluid intelligence. Human Brain Mapping, 45(3), e26591. 10.1002/hbm.26591

Piche, E., Chorin, F., Gerus, P., Jaafar, A., Guerin, O., & Zory, R. (2023). Effects of age, sex, frailty and falls on cognitive and motor performance during dual-task walking in older adults. Experimental Gerontology, 171, 112022. 10.1016/j.exger.2022.112022

Plummer-D’Amato, P., Brancato, B., Dantowitz, M., Birken, S., Bonke, C., & Furey, E. (2012). Effects of gait and cognitive task difficulty on cognitive-motor interference in aging. Journal of Aging Research, 2012, 583894. 10.1155/2012/583894

Promjunyakul, N., Schmit, B. D., & Schindler-Ivens, S. M. (2015). A novel fMRI paradigm suggests that pedaling-related brain activation is altered after stroke. Frontiers in Human Neuroscience, 09. 10.3389/fnhum.2015.00324

Reinhardt, J., Rus-Oswald, O. G., Bürki, C. N., Bridenbaugh, S. A., Krumm, S., Michels, L., Stippich, C., Kressig, R. W., & Blatow, M. (2020). Neural Correlates of Stepping in Healthy Elderly: Parietal and Prefrontal Cortex Activation Reflects Cognitive-Motor Interference Effects. Frontiers in Human Neuroscience, 14, 566735. 10.3389/fnhum.2020.566735

Riccardi, N., Teghipco, A., Newman-Norlund, S., Newman-Norlund, R., Rangus, I., Rorden, C., Fridriksson, J., & Bonilha, L. (2025). Distinct brain age gradients across the adult lifespan reflect diverse neurobiological hierarchies. Communications Biology, 8(1), 802. 10.1038/s42003-025-08228-z

Rohart, F., Gautier, B., Singh, A., & Lê Cao, K.-A. (2017). mixOmics: An R package for ’omics feature selection and multiple data integration. PLoS Computational Biology, 13(11), e1005752. 10.1371/journal.pcbi.1005752

Schäfer, J., & Strimmer, K. (2005). A shrinkage approach to large-scale covariance matrix estimation and implications for functional genomics. Statistical Applications in Genetics and Molecular Biology, 4, Article32. 10.2202/1544-6115.1175

Seidler, R. D., Bernard, J. A., Burutolu, T. B., Fling, B. W., Gordon, M. T., Gwin, J. T., Kwak, Y., & Lipps, D. B. (2010). Motor control and aging: Links to age-related brain structural, functional, and biochemical effects. Neurosci Biobehav Rev, 34(5), 721–733. 10.1016/j.neubiorev.2009.10.005

Song, J., Birn, R. M., Boly, M., Meier, T. B., Nair, V. A., Meyerand, M. E., & Prabhakaran, V. (2014). Age-Related Reorganizational Changes in Modularity and Functional Connectivity of Human Brain Networks. Brain Connectivity, 4(9), 662–676. 10.1089/brain.2014.0286

St George, R. J., Jayakody, O., Healey, R., Breslin, M., Hinder, M. R., & Callisaya, M. L. (2022). Cognitive inhibition tasks interfere with dual-task walking and increase prefrontal cortical activity more than working memory tasks in young and older adults. Gait & Posture, 95, 186–191. 10.1016/j.gaitpost.2022.04.021

Stojan, R., Mack, M., Bock, O., & Voelcker-Rehage, C. (2023). Inefficient frontal and parietal brain activation during dual-task walking in a virtual environment in older adults. NeuroImage, 273, 120070. 10.1016/j.neuroimage.2023.120070

Tenenhaus, M., Tenenhaus, A., & Groenen, P. J. F. (2017). Regularized Generalized Canonical Correlation Analysis: A Framework for Sequential Multiblock Component Methods. Psychometrika, 82(3), 737–777. 10.1007/s11336-017-9573-x

Tomasi, D., & Volkow, N. D. (2012). Aging and functional brain networks. Molecular Psychiatry, 17(5), 471, 549–558. 10.1038/mp.2011.81

Udina, C., Ayers, E., Inzitari, M., & Verghese, J. (2021). Walking While Talking and Prefrontal Oxygenation in Motoric Cognitive Risk Syndrome: Clinical and Pathophysiological Aspects. Journal of Alzheimer’s Disease, 84(4), 1585–1596. 10.3233/JAD-210239

Vinehout, K., Schmit, B. D., & Schindler-Ivens, S. (2019). Lower Limb Task-Based Functional Connectivity Is Altered in Stroke. Brain Connectivity, 9(4), 365–377. 10.1089/brain.2018.0640

Wang, B., LeBel, A., & D’Mello, A. M. (2025). Ignoring the cerebellum is hindering progress in neuroscience. Trends in Cognitive Sciences, 29(4), 318–330. 10.1016/j.tics.2025.01.004

Wang, X., Huang, C.-C., Tsai, S.-J., Lin, C.-P., & Cai, Q. (2024). The aging trajectories of brain functional hierarchy and its impact on cognition across the adult lifespan. Frontiers in Aging Neuroscience, 16, 1331574. 10.3389/fnagi.2024.1331574

Ward, N. S., & Frackowiak, R. S. J. (2003). Age-related changes in the neural correlates of motor performance. Brain: A Journal of Neurology, 126(Pt 4), 873–888. 10.1093/brain/awg071

Welham, Z., Déjean, S., & Lê Cao, K.-A. (2023). Multivariate Analysis with the R Package mixOmics. *Methods in Molecular Biology (Clifton*, N.J.), 2426, 333–359. 10.1007/978-1-0716-1967-4_15

Whitfield-Gabrieli, S., & Nieto-Castanon, A. (2012). Conn: A functional connectivity toolbox for correlated and anticorrelated brain networks. Brain Connect, 2(3), 125–141. 10.1089/brain.2012.0073

Wilson, R. S., Wang, T., Yu, L., Bennett, D. A., & Boyle, P. A. (2020). Normative Cognitive Decline in Old Age. Annals of Neurology, 87(6), 816–829. 10.1002/ana.25711

Xu, J., Moeller, S., Auerbach, E. J., Strupp, J., Smith, S. M., Feinberg, D. A., Yacoub, E., & Uğurbil, K. (2013). Evaluation of slice accelerations using multiband echo planar imaging at 3 T. NeuroImage, 83, 991–1001. 10.1016/j.neuroimage.2013.07.055

Yang, Q., Tian, C., Tseng, B., Zhang, B., Huang, S., Jin, S., & Mao, J. (2020). Gait Change in Dual Task as a Behavioral Marker to Detect Mild Cognitive Impairment in Elderly Persons: A Systematic Review and Meta-analysis. Archives of Physical Medicine and Rehabilitation, 101(10), 1813– 1821. 10.1016/j.apmr.2020.05.020

Yuan, J., Blumen, H. M., Verghese, J., & Holtzer, R. (2015). Functional connectivity associated with gait velocity during walking and walking-while-talking in aging: A resting-state fMRI study. Human Brain Mapping, 36(4), 1484–1493. 10.1002/hbm.22717

Zimmermann, P., & Fimm, B. (2002). A test battery for attentional performance. In M. Leclercq & P. Zimmermann (Eds), Applied Neuropsychology of Attention (pp. 110–151). Psychology Press. 10.4324/9780203307014-12

