## Supplemental Materials for "Domain-Specific Functional Network Adaptations Supporting Dual-Task Performance in Older Adults"

Address:

Department of Psychology, Building A07,

26111 Oldenburg,

Germany

**Supplementary Material A:** Preprocess through *fMRIPrep*

Results included in this manuscript come from preprocessing performed using fMRIPrep 24.1.1 (Esteban et al. (2019); Esteban et al. (2018); RRID:SCR_016216), which is based on Nipype 1.8.6 (K. Gorgolewski et al. (2011); K. J. Gorgolewski et al. (2018); RRID:SCR_002502).

Preprocessing of B_0_ inhomogeneity mappings

A total of 2 fieldmaps were found available within the input BIDS structure for this particular subject. A B_0_ nonuniformity map (or fieldmap) was estimated from the phase-drift map(s) measure with two consecutive GRE (gradient-recalled echo) acquisitions. The corresponding phase-map(s) were phase-unwrapped with prelude (FSL None).

Anatomical data preprocessing

A total of 1 T1-weighted (T1w) images were found within the input BIDS dataset. The T1w image was corrected for intensity non-uniformity (INU) with N4BiasFieldCorrection (Tustison et al. 2010), distributed with ANTs 2.5.3 (Avants et al. 2008, RRID:SCR_004757), and used as T1w-reference throughout the workflow. The T1w-reference was then skull-stripped with a Nipype implementation of the antsBrainExtraction.sh workflow (from ANTs), using OASIS30ANTs as target template. Brain tissue segmentation of cerebrospinal fluid (CSF), white-matter (WM) and gray-matter (GM) was performed on the brain-extracted T1w using fast (FSL (version unknown), RRID:SCR_002823, Zhang, Brady, and Smith 2001). Brain surfaces were reconstructed using recon-all (FreeSurfer 7.3.2, RRID:SCR_001847, Dale, Fischl, and Sereno 1999), and the brain mask estimated previously was refined with a custom variation of the method to reconcile ANTs-derived and FreeSurfer-derived segmentations of the cortical gray-matter of Mindboggle (RRID:SCR_002438, Klein et al. 2017). A T2-weighted image was used to improve pial surface refinement. Brain surfaces were reconstructed using recon-all (FreeSurfer 7.3.2, RRID:SCR_001847, Dale, Fischl, and Sereno 1999), and the brain mask estimated previously was refined with a custom variation of the method to reconcile ANTs-derived and FreeSurfer-derived segmentations of the cortical gray-matter of Mindboggle (RRID:SCR_002438, Klein et al. 2017). Volume-based spatial normalization to one standard space (MNI152NLin2009cAsym) was performed through nonlinear registration with antsRegistration (ANTs 2.5.3), using brain-extracted versions of both T1w reference and the T1w template. The following template was were selected for spatial normalization and accessed with TemplateFlow (24.2.0, Ciric et al. 2022): ICBM 152 Nonlinear Asymmetrical template version 2009c [Fonov et al. (2009), RRID:SCR_008796; TemplateFlow ID: MNI152NLin2009cAsym].

Functional data preprocessing

For each of the 3 BOLD runs found per subject (across all tasks and sessions), the following preprocessing was performed. First, a reference volume was generated, using a custom methodology of fMRIPrep, for use in head motion correction. Head-motion parameters with respect to the BOLD reference (transformation matrices, and six corresponding rotation and translation parameters) are estimated before any spatiotemporal filtering using mcflirt (FSL , Jenkinson et al. 2002). The BOLD reference was then co-registered to the T1w reference using bbregister (FreeSurfer) which implements boundary-based registration (Greve and Fischl 2009). Co-registration was configured with six degrees of freedom. The aligned T2w image was used for initial co-registration.Several confounding time-series were calculated based on the preprocessed BOLD: framewise displacement (FD), DVARS and three region-wise global signals. FD was computed using two formulations following Power (absolute sum of relative motions, Power et al. (2014)) and Jenkinson (relative root mean square displacement between affines, Jenkinson et al. (2002)). FD and DVARS are calculated for each functional run, both using their implementations in Nipype (following the definitions by Power et al. 2014). The three global signals are extracted within the CSF, the WM, and the whole-brain masks. Additionally, a set of physiological regressors were extracted to allow for component-based noise correction (CompCor, Behzadi et al. 2007). Principal components are estimated after high-pass filtering the preprocessed BOLD time-series (using a discrete cosine filter with 128s cut-off) for the two CompCor variants: temporal (tCompCor) and anatomical (aCompCor). tCompCor components are then calculated from the top 2% variable voxels within the brain mask. For aCompCor, three probabilistic masks (CSF, WM and combined CSF+WM) are generated in anatomical space. The implementation differs from that of Behzadi et al. in that instead of eroding the masks by 2 pixels on BOLD space, a mask of pixels that likely contain a volume fraction of GM is subtracted from the aCompCor masks. This mask is obtained by dilating a GM mask extracted from the FreeSurfer’s aseg segmentation, and it ensures components are not extracted from voxels containing a minimal fraction of GM. Finally, these masks are resampled into BOLD space and binarized by thresholding at 0.99 (as in the original implementation). Components are also calculated separately within the WM and CSF masks. For each CompCor decomposition, the k components with the largest singular values are retained, such that the retained components’ time series are sufficient to explain 50 percent of variance across the nuisance mask (CSF, WM, combined, or temporal). The remaining components are dropped from consideration. The head-motion estimates calculated in the correction step were also placed within the corresponding confounds file. The confound time series derived from head motion estimates and global signals were expanded with the inclusion of temporal derivatives and quadratic terms for each (Satterthwaite et al. 2013). Frames that exceeded a threshold of 0.5 mm FD or 1.5 standardized DVARS were annotated as motion outliers. Additional nuisance timeseries are calculated by means of principal components analysis of the signal found within a thin band (crown) of voxels around the edge of the brain, as proposed by (Patriat, Reynolds, and Birn 2017). All resamplings can be performed with a single interpolation step by composing all the pertinent transformations (i.e. head-motion transform matrices, susceptibility distortion correction when available, and co-registrations to anatomical and output spaces). Gridded (volumetric) resamplings were performed using nitransforms, configured with cubic B-spline interpolation.

Many internal operations of fMRIPrep use Nilearn 0.10.4 (Abraham et al. 2014, RRID:SCR_001362), mostly within the functional processing workflow. For more details of the pipeline, see [the section corresponding to workflows in fMRIPrep’s documentation](https://fmriprep.readthedocs.io/en/latest/workflows.html).

##### Copyright Waiver

The above boilerplate text was automatically generated by fMRIPrep with the express intention that users should copy and paste this text into their manuscripts unchanged. It is released under the [CC0](https://creativecommons.org/publicdomain/zero/1.0/) license.

**S****upplementary Figure S1.** Group-level whole-brain activation patterns for the cognitive single Go, cognitive single NoGo, single motor and dual-task contrast across all subjects. Significant activations were identified using a family-wise error (FWE)–corrected threshold of *p* < 0.05. The dual-task condition elicited the strongest and most extensive activations, encompassing the precentral gyrus, supplementary motor area (SMA), insular cortex, basal ganglia, and large regions of the cerebellum (lobules I–IV).

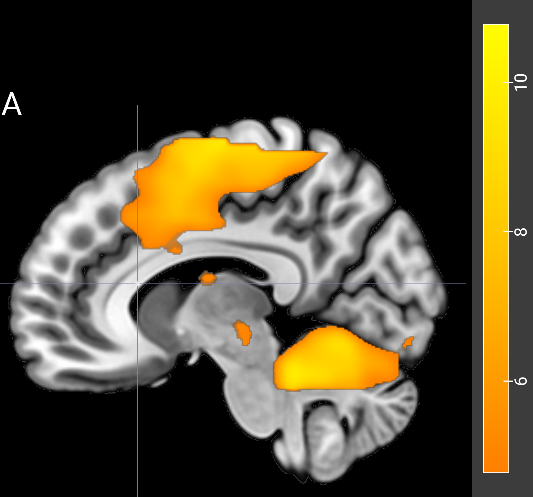

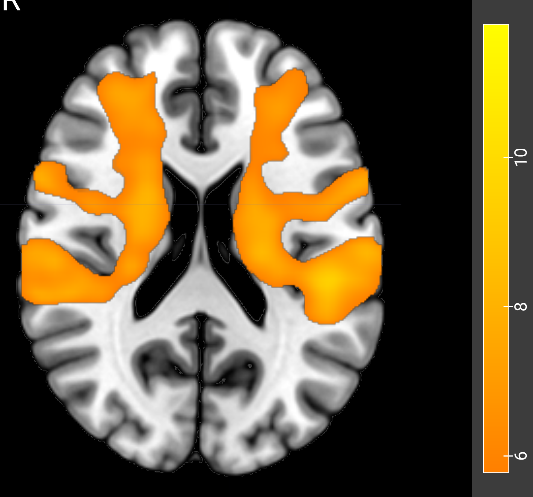

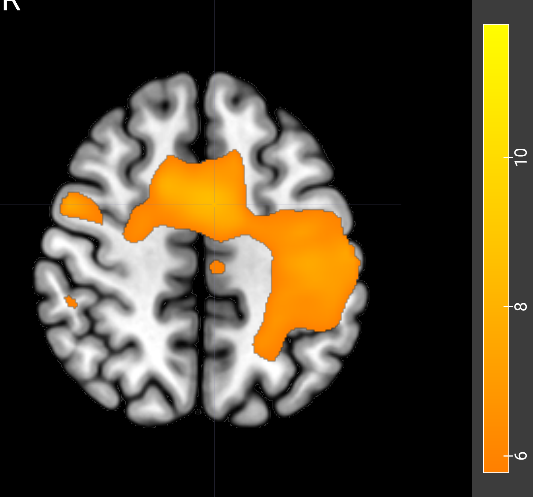

z50

z20

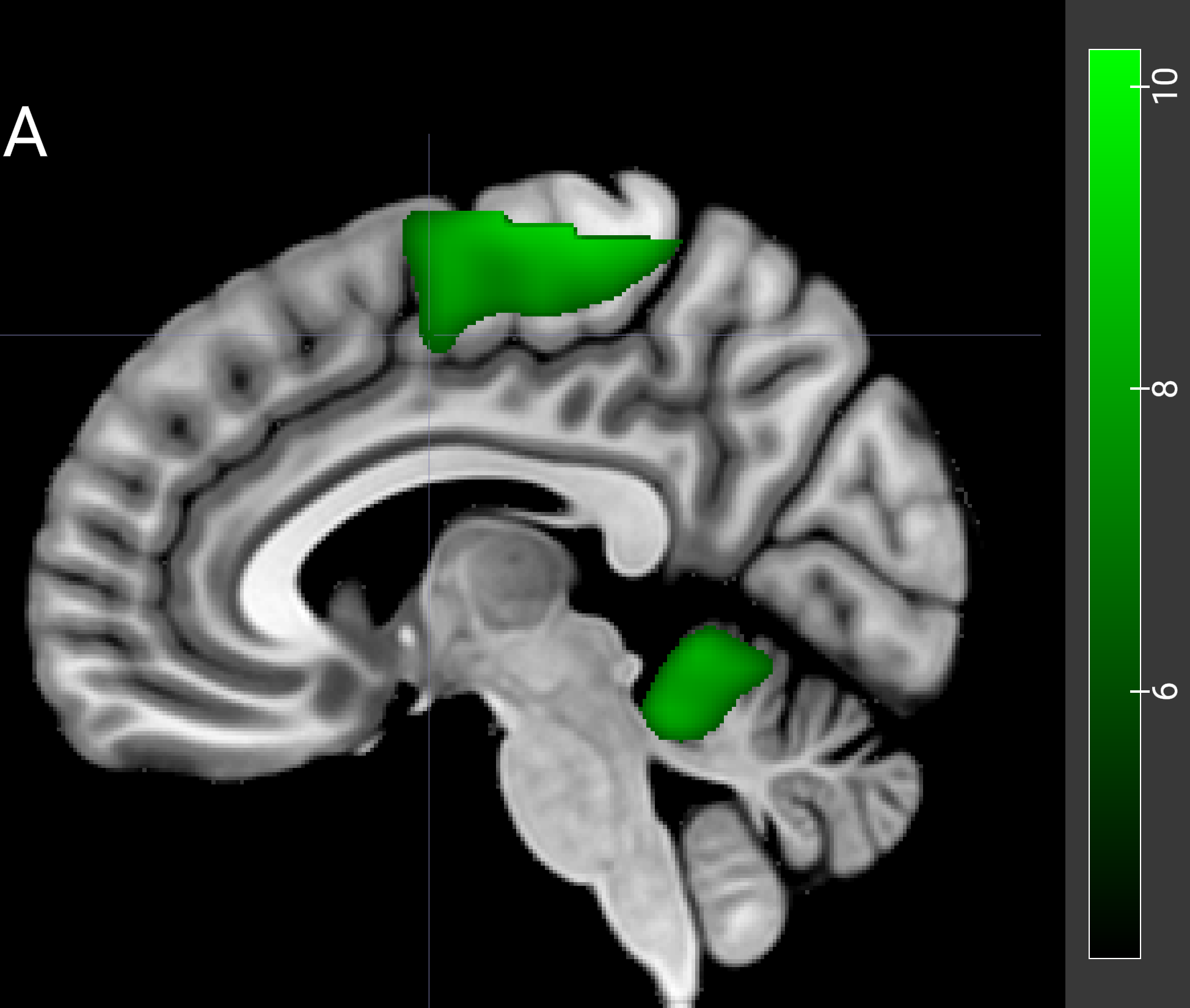

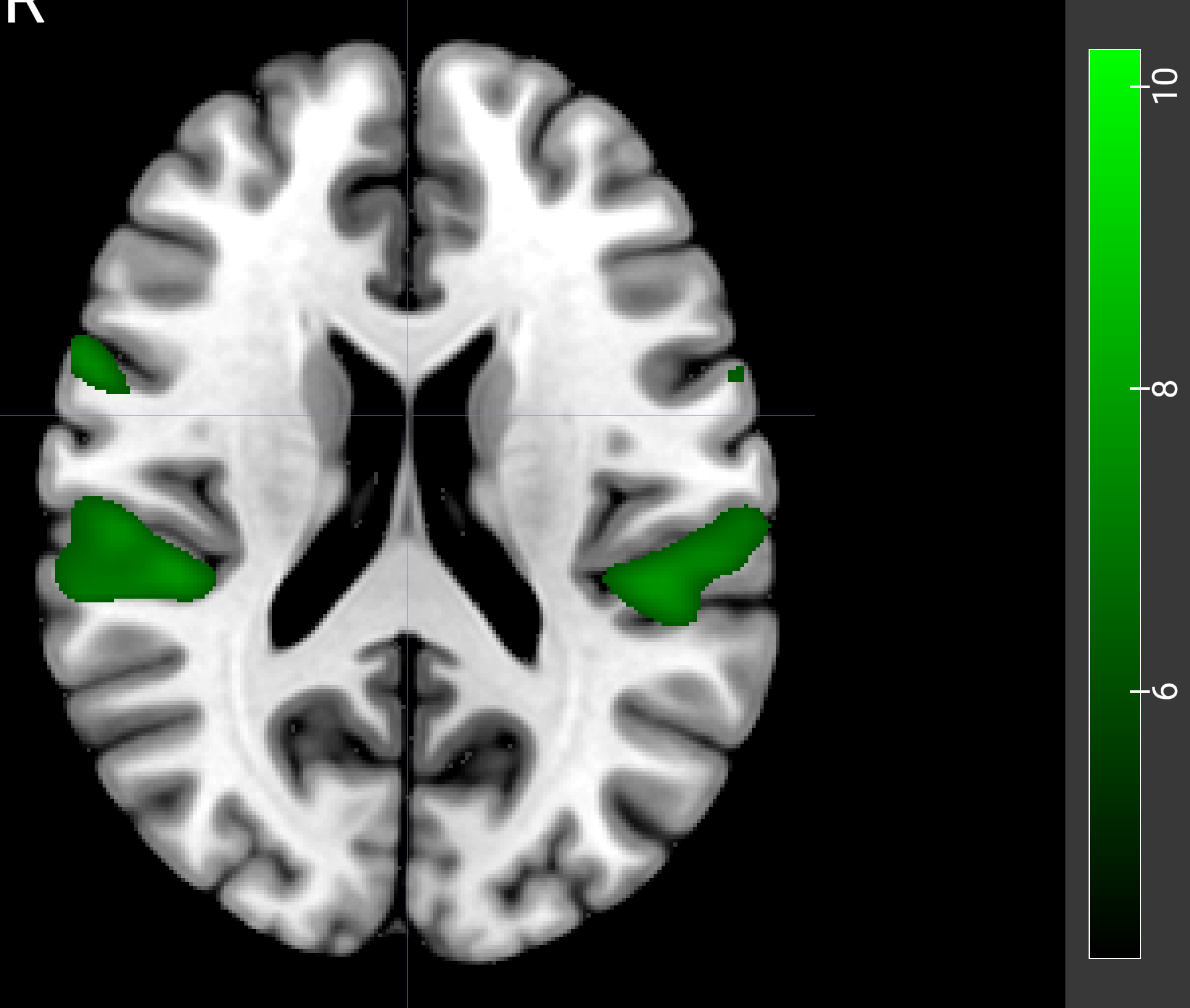

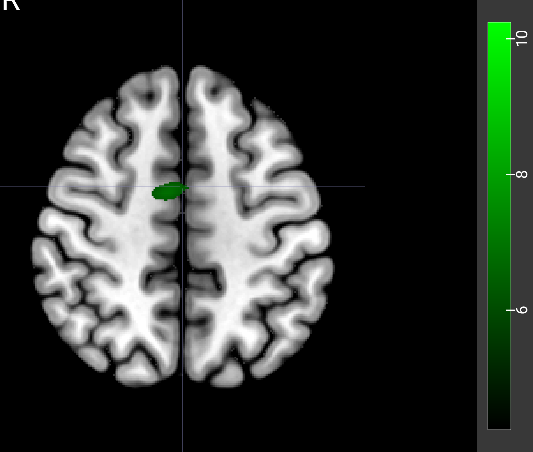

sagittal

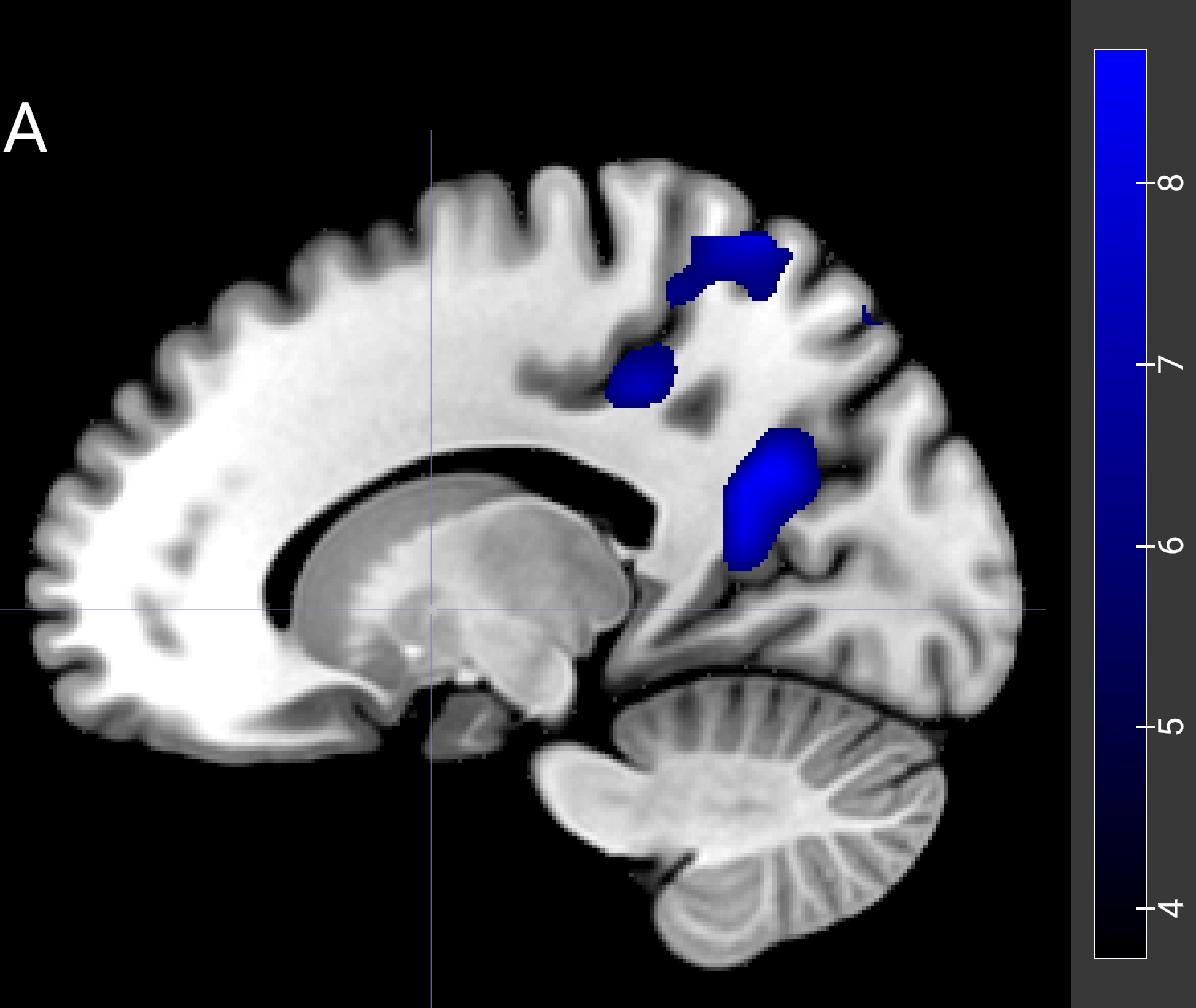

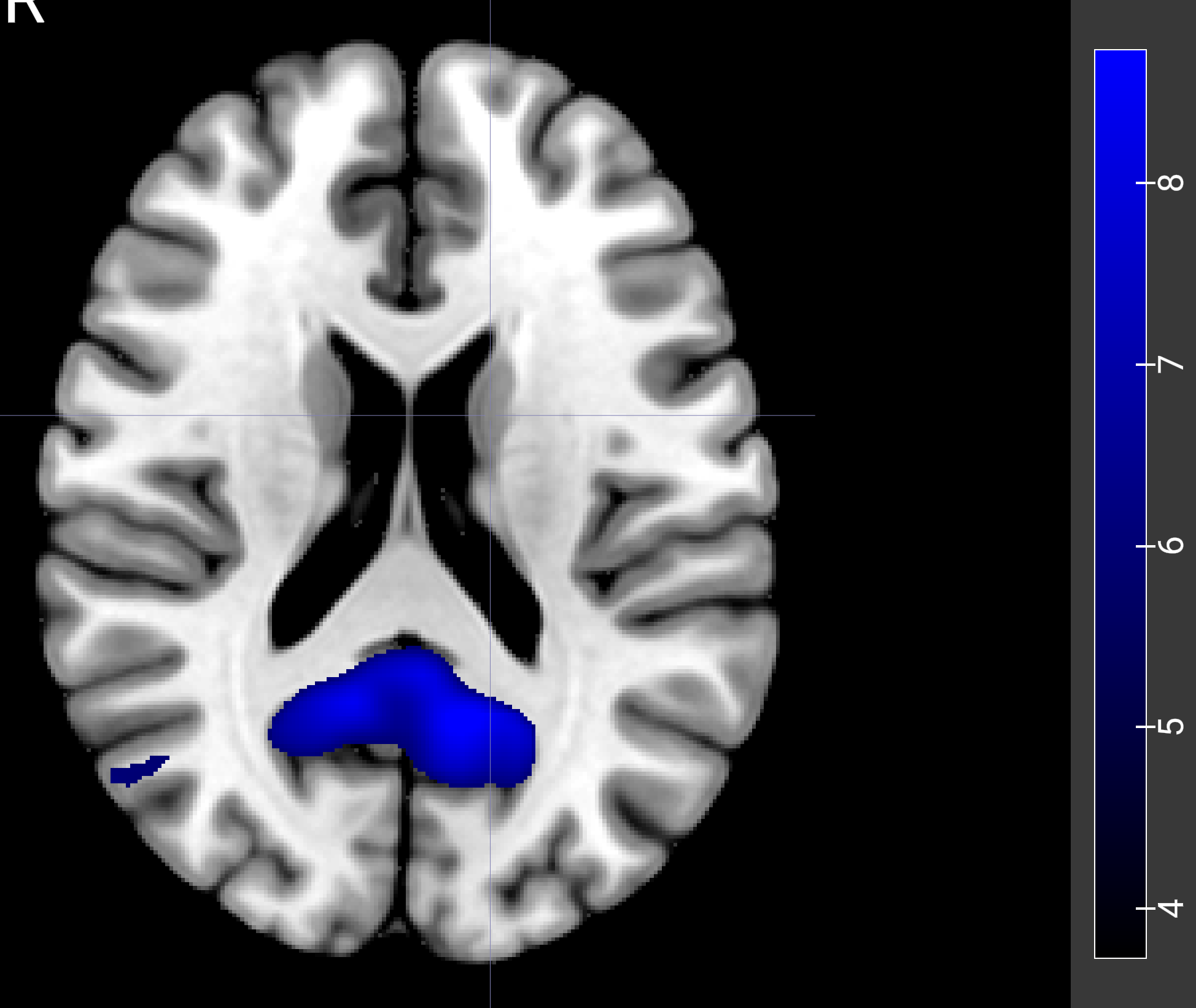

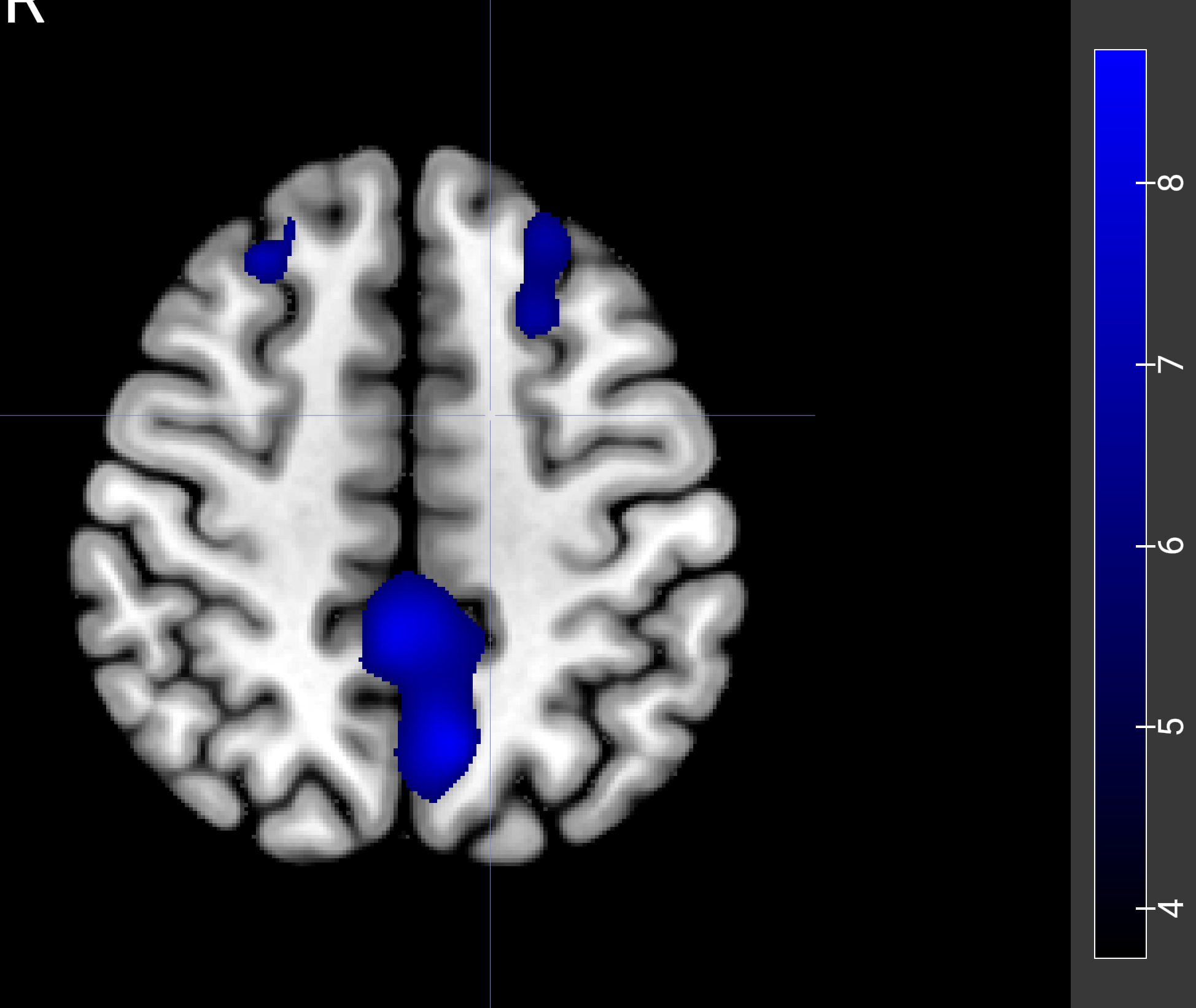

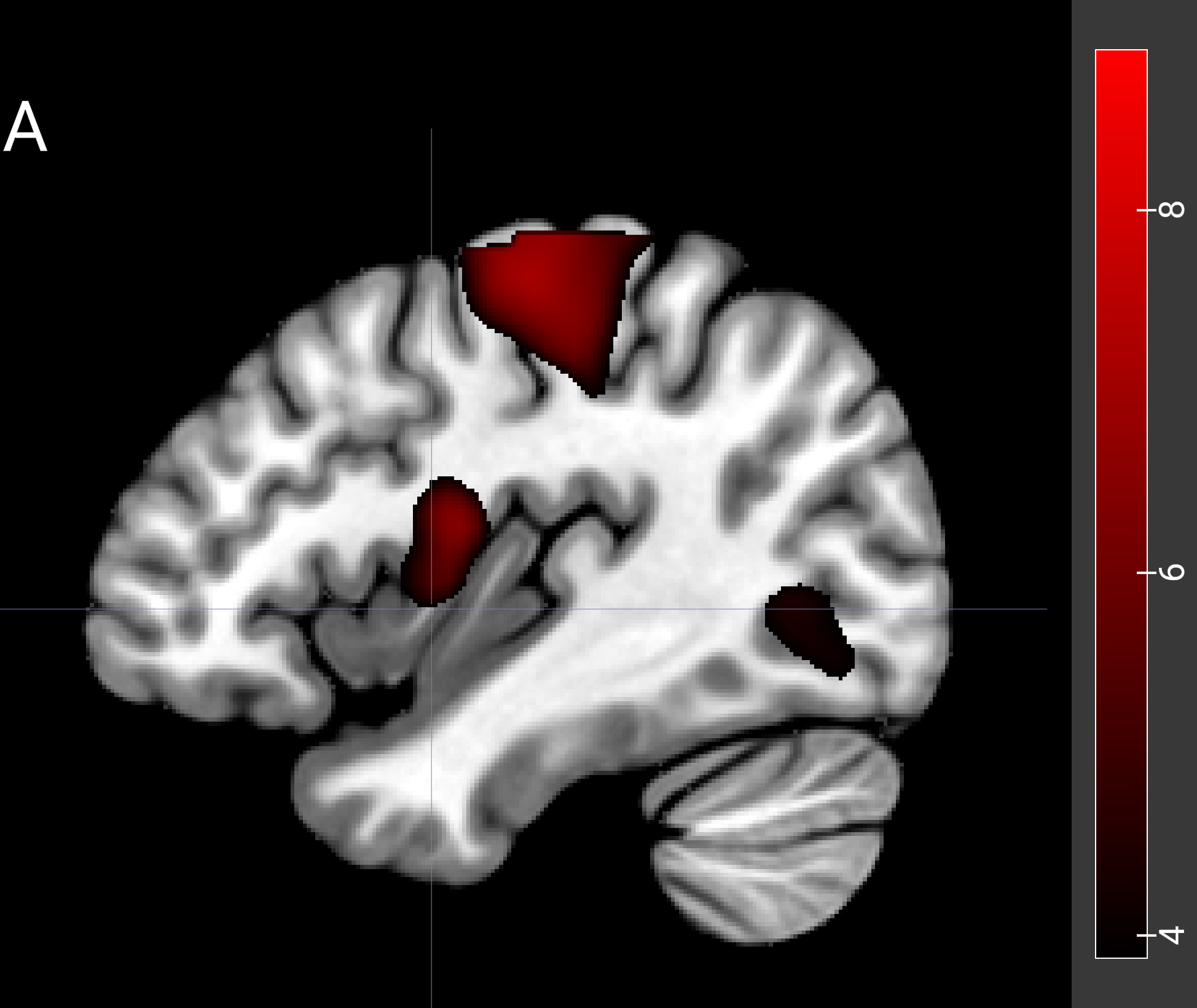

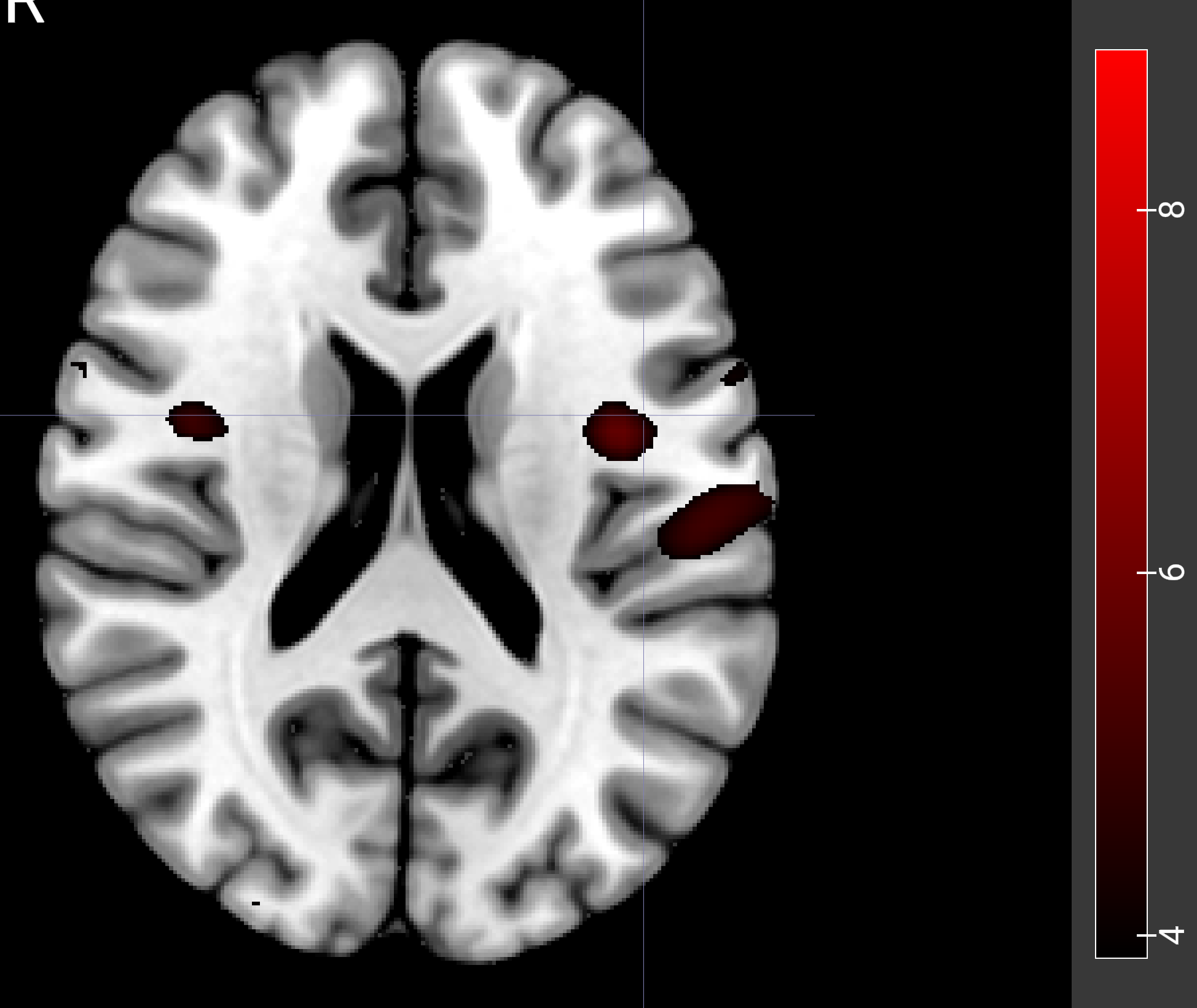

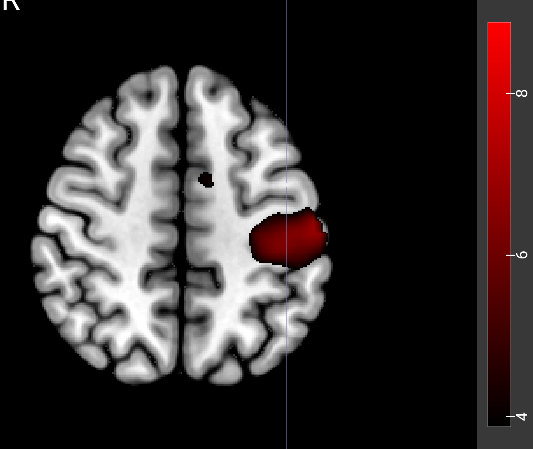

R

R

Dual task

Single motor

Single Go

-Single NoGo

**Supplementary Table S1.** Regions of interest (ROIs) identified from the dual-task contrasts. ROIs prefixed with C_ROI_ were derived from the cognitive-domain contrast (Dual Go > Single Motor), and ROIs prefixed with M_ROI_ were derived from the motor-domain contrast (Dual Go > Single Cognitive Go). ROIs ending with *_l* and *_r* indicate the left and right hemispheres, respectively. Cluster sizes are reported in voxels. Abbreviations of ROIs are provided at the end of table.

### Cognitive Domain (C_ROI)

#### Frontal regions

| ROI | Cluster size (voxels) |
| --- | --- |
| C_ROI_FP_l | 141 |
| C_ROI_FP_r | 93 |
| C_ROI_IFGoper_l | 35 |
| C_ROI_MidFG_l | 342 |
| C_ROI_MidFG_r | 166 |
| C_ROI_SFG_l | 292 |
| C_ROI_SFG_r | 225 |
| C_ROI_SMA_l | 175 |
| C_ROI_SMA_r | 85 |
| C_ROI_PreCG_l | 783 |
| C_ROI_PreCG_r | 213 |

#### Parietal regions

| ROI | Cluster size (voxels) |
| --- | --- |
| C_ROI_AG_l | 31 |
| C_ROI_AG_r | 23 |
| C_ROI_OP_l | 106 |
| C_ROI_OP_r | 75 |
| C_ROI_PaCiG_l | 62 |
| C_ROI_PaCiG_r | 48 |
| C_ROI_PostCG_l | 681 |
| C_ROI_SPL_l | 457 |
| C_ROI_SPL_r | 200 |
| C_ROI_aSMG_l | 122 |
| C_ROI_aSMG_r | 31 |
| C_ROI_pSMG_l | 143 |
| C_ROI_pSMG_r | 78 |
| C_ROI_sLOC_l | 1087 |
| C_ROI_sLOC_r | 971 |

#### Temporal/Occipital regions

| ROI | Cluster size (voxels) |
| --- | --- |
| C_ROI_LG_l | 195 |
| C_ROI_LG_r | 223 |
| C_ROI_OFusG_l | 250 |
| C_ROI_OFusG_r | 262 |
| C_ROI_TOFusC_l | 32 |
| C_ROI_TOFusC_r | 79 |
| C_ROI_iLOC_l | 178 |
| C_ROI_iLOC_r | 59 |
| C_ROI_toITG_l | 22 |

#### Subcortical regions

| ROI | Cluster size (voxels) |
| --- | --- |
| C_ROI_Caudate_r | 34 |
| C_ROI_Putamen_l | 66 |

#### Cerebellar / Midline regions

| ROI | Cluster size (voxels) |
| --- | --- |
| C_ROI_Cereb1_l | 303 |
| C_ROI_Cereb1_r | 181 |
| C_ROI_Cereb2_l | 50 |
| C_ROI_Cereb45_r | 134 |
| C_ROI_Cereb6_l | 216 |
| C_ROI_Cereb6_r | 551 |
| C_ROI_Ver4_5 | 65 |
| C_ROI_Ver6 | 158 |
| C_ROI_Ver7 | 72 |

### **Motor Domain (M_ROI)**

#### Frontal regions

| ROI | Cluster size (voxels) |
| --- | --- |
| M_ROI_FP_l | 616 |
| M_ROI_FP_r | 788 |
| M_ROI_IFGoper_l | 152 |
| M_ROI_IFGoper_r | 199 |
| M_ROI_MidFG_l | 606 |
| M_ROI_MidFG_r | 623 |
| M_ROI_PreCG_l | 1320 |
| M_ROI_PreCG_r | 1066 |
| M_ROI_SFG_l | 543 |
| M_ROI_SFG_r | 664 |
| M_ROI_SMA_l | 315 |
| M_ROI_SMA_r | 366 |
| M_ROI_FO_l | 67 |

#### Parietal regions

| ROI | Cluster size (voxels) |
| --- | --- |
| M_ROI_AG_l | 101 |
| M_ROI_AG_r | 73 |
| M_ROI_PO_l | 253 |
| M_ROI_PO_r | 241 |
| M_ROI_PT_l | 129 |
| M_ROI_PT_r | 162 |
| M_ROI_PaCiG_l | 87 |
| M_ROI_PaCiG_r | 105 |
| M_ROI_PostCG_l | 747 |
| M_ROI_PostCG_r | 545 |
| M_ROI_Precuneus | 751 |
| M_ROI_SPL_l | 667 |
| M_ROI_SPL_r | 650 |
| M_ROI_aSMG_l | 398 |
| M_ROI_aSMG_r | 356 |
| M_ROI_pSMG_l | 370 |
| M_ROI_pSMG_r | 377 |

#### Temporal/Occipital regions

| ROI | Cluster size (voxels) |
| --- | --- |
| M_ROI_LG_l | 106 |
| M_ROI_LG_r | 97 |
| M_ROI_OFusG_l | 54 |
| M_ROI_TOFusC_l | 34 |
| M_ROI_iLOC_l | 33 |
| M_ROI_iLOC_r | 36 |
| M_ROI_pSTG_l | 22 |
| M_ROI_sLOC_l | 866 |
| M_ROI_sLOC_r | 647 |
| M_ROI_toMTG_l | 49 |

#### Subcortical regions

| ROI | Cluster size (voxels) |
| --- | --- |
| M_ROI_Caudate_l | 53 |
| M_ROI_Caudate_r | 78 |
| M_ROI_Putamen_l | 110 |
| M_ROI_Putamen_r | 120 |
| M_ROI_Pallidum_l | 50 |
| M_ROI_Pallidum_r | 42 |
| M_ROI_Thalamus_l | 236 |
| M_ROI_Thalamus_r | 225 |

**Abbreviations of ROIs**

FP r (Frontal Pole Right)

FP l (Frontal Pole Left)

IC r (Insular Cortex Right)

IC l (Insular Cortex Left)

SFG r (Superior Frontal Gyrus Right)

SFG l (Superior Frontal Gyrus Left)

MidFG r (Middle Frontal Gyrus Right)

MidFG l (Middle Frontal Gyrus Left)

IFG tri r (Inferior Frontal Gyrus, pars triangularis Right)

IFG tri l (Inferior Frontal Gyrus, pars triangularis Left)

IFG oper r (Inferior Frontal Gyrus, pars opercularis Right)

IFG oper l (Inferior Frontal Gyrus, pars opercularis Left)

PreCG r (Precentral Gyrus Right)

PreCG l (Precentral Gyrus Left)

TP r (Temporal Pole Right)

TP l (Temporal Pole Left)

aSTG r (Superior Temporal Gyrus, anterior division Right)

aSTG l (Superior Temporal Gyrus, anterior division Left)

pSTG r (Superior Temporal Gyrus, posterior division Right)

pSTG l (Superior Temporal Gyrus, posterior division Left)

aMTG r (Middle Temporal Gyrus, anterior division Right)

aMTG l (Middle Temporal Gyrus, anterior division Left)

pMTG r (Middle Temporal Gyrus, posterior division Right)

pMTG l (Middle Temporal Gyrus, posterior division Left)

toMTG r (Middle Temporal Gyrus, temporooccipital part Right)

toMTG l (Middle Temporal Gyrus, temporooccipital part Left)

aITG r (Inferior Temporal Gyrus, anterior division Right)

aITG l (Inferior Temporal Gyrus, anterior division Left)

pITG r (Inferior Temporal Gyrus, posterior division Right)

pITG l (Inferior Temporal Gyrus, posterior division Left)

toITG r (Inferior Temporal Gyrus, temporooccipital part Right)

toITG l (Inferior Temporal Gyrus, temporooccipital part Left)

PostCG r (Postcentral Gyrus Right)

PostCG l (Postcentral Gyrus Left)

SPL r (Superior Parietal Lobule Right)

SPL l (Superior Parietal Lobule Left)

aSMG r (Supramarginal Gyrus, anterior division Right)

aSMG l (Supramarginal Gyrus, anterior division Left)

pSMG r (Supramarginal Gyrus, posterior division Right)

pSMG l (Supramarginal Gyrus, posterior division Left)

AG r (Angular Gyrus Right)

AG l (Angular Gyrus Left)

sLOC r (Lateral Occipital Cortex, superior division Right)

sLOC l (Lateral Occipital Cortex, superior division Left)

iLOC r (Lateral Occipital Cortex, inferior division Right)

iLOC l (Lateral Occipital Cortex, inferior division Left)

ICC r (Intracalcarine Cortex Right)

ICC l (Intracalcarine Cortex Left)

MedFC (Frontal Medial Cortex)

SMA r (Juxtapositional Lobule Cortex -formerly Supplementary Motor Cortex- Right)

SMA L(Juxtapositional Lobule Cortex -formerly Supplementary Motor Cortex- Left)

SubCalC (Subcallosal Cortex)

PaCiG r (Paracingulate Gyrus Right)

PaCiG l (Paracingulate Gyrus Left)

AC (Cingulate Gyrus, anterior division)

PC (Cingulate Gyrus, posterior division)

Precuneous (Precuneous Cortex)

Cuneal r (Cuneal Cortex Right)

Cuneal l (Cuneal Cortex Left)

FOrb r (Frontal Orbital Cortex Right)

FOrb l (Frontal Orbital Cortex Left)

aPaHC r (Parahippocampal Gyrus, anterior division Right)

aPaHC l (Parahippocampal Gyrus, anterior division Left)

pPaHC r (Parahippocampal Gyrus, posterior division Right)

pPaHC l (Parahippocampal Gyrus, posterior division Left)

LG r (Lingual Gyrus Right)

LG l (Lingual Gyrus Left)

aTFusC r (Temporal Fusiform Cortex, anterior division Right)

aTFusC l (Temporal Fusiform Cortex, anterior division Left)

pTFusC r (Temporal Fusiform Cortex, posterior division Right)

pTFusC l (Temporal Fusiform Cortex, posterior division Left)

TOFusC r (Temporal Occipital Fusiform Cortex Right)

TOFusC l (Temporal Occipital Fusiform Cortex Left)

OFusG r (Occipital Fusiform Gyrus Right)

OFusG l (Occipital Fusiform Gyrus Left)

FO r (Frontal Operculum Cortex Right)

FO l (Frontal Operculum Cortex Left)

CO r (Central Opercular Cortex Right)

CO l (Central Opercular Cortex Left)

PO r (Parietal Operculum Cortex Right)

PO l (Parietal Operculum Cortex Left)

PP r (Planum Polare Right)

PP l (Planum Polare Left)

HG r (Heschl's Gyrus Right)

HG l (Heschl's Gyrus Left)

PT r (Planum Temporale Right)

PT l (Planum Temporale Left)

SCC r (Supracalcarine Cortex Right)

SCC l (Supracalcarine Cortex Left)

OP r (Occipital Pole Right)

OP l (Occipital Pole Left)

Thalamus r

Thalamus l

Caudate r

Caudate l

Putamen r

Putamen l

Pallidum r

Pallidum l

Hippocampus r

Hippocampus l

Amygdala r

Amygdala l

Accumbens r

Accumbens l

Brain-Stem

Cereb1 l (Cerebelum Crus1 Left)

Cereb1 r (Cerebelum Crus1 Right)

Cereb2 l (Cerebelum Crus2 Left)

Cereb2 r (Cerebelum Crus2 Right)

Cereb3 l (Cerebelum 3 Left)

Cereb3 r (Cerebelum 3 Right)

Cereb45 l (Cerebelum 4 5 Left)

Cereb45 r (Cerebelum 4 5 Right)

Cereb6 l (Cerebelum 6 Left)

Cereb6 r (Cerebelum 6 Right)

Cereb7 l (Cerebelum 7b Left)

Cereb7 r (Cerebelum 7b Right)

Cereb8 l (Cerebelum 8 Left)

Cereb8 r (Cerebelum 8 Right)

Cereb9 l (Cerebelum 9 Left)

Cereb9 r (Cerebelum 9 Right)

Cereb10 l (Cerebelum 10 Left)

Cereb10 r (Cerebelum 10 Right)

Ver12 (Vermis 1 2)

Ver3 (Vermis 3)

Ver45 (Vermis 4 5)

Ver6 (Vermis 6)

Ver7 (Vermis 7)

Ver8 (Vermis 8)

Ver9 (Vermis 9)

Ver10 (Vermis 10)

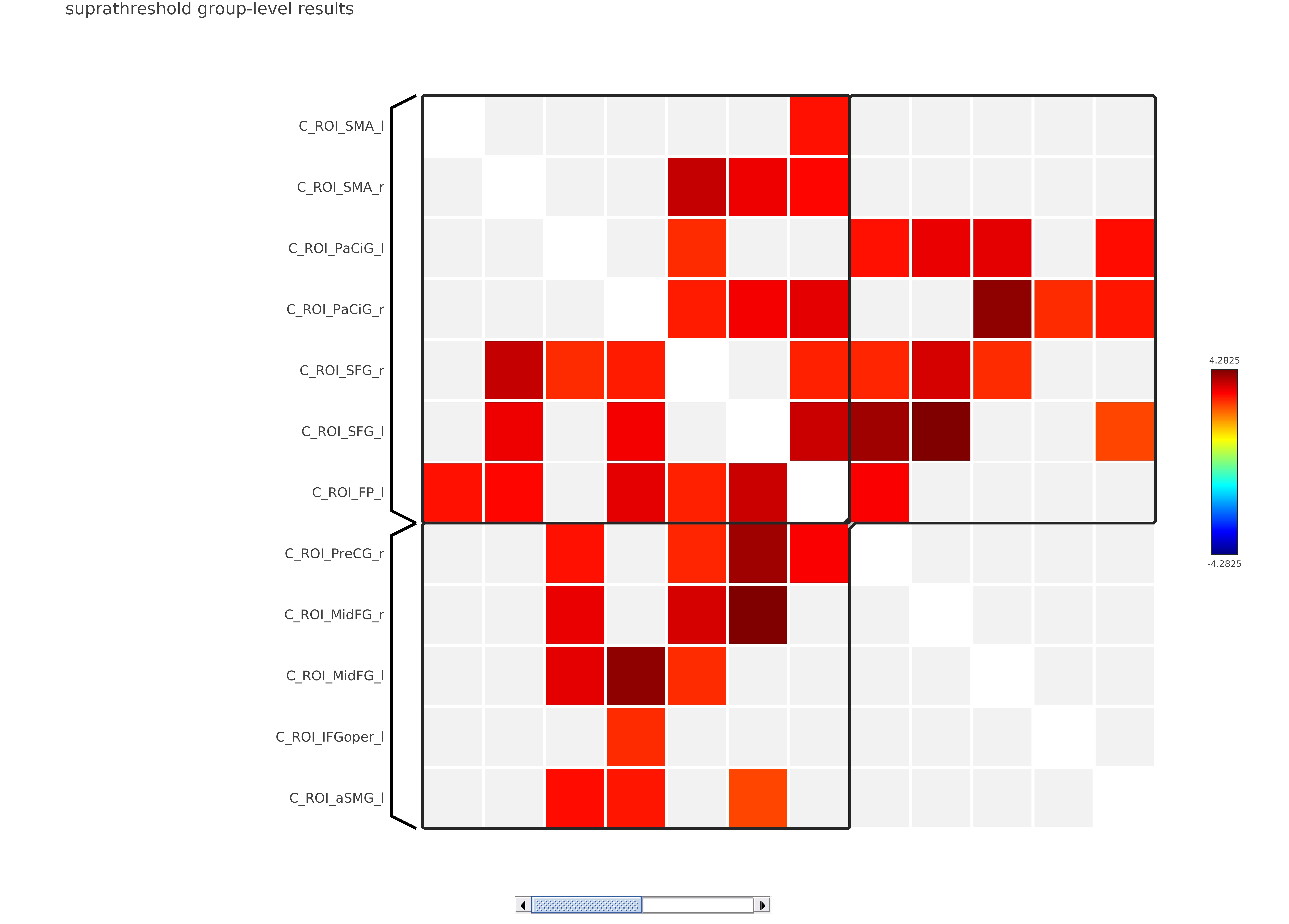

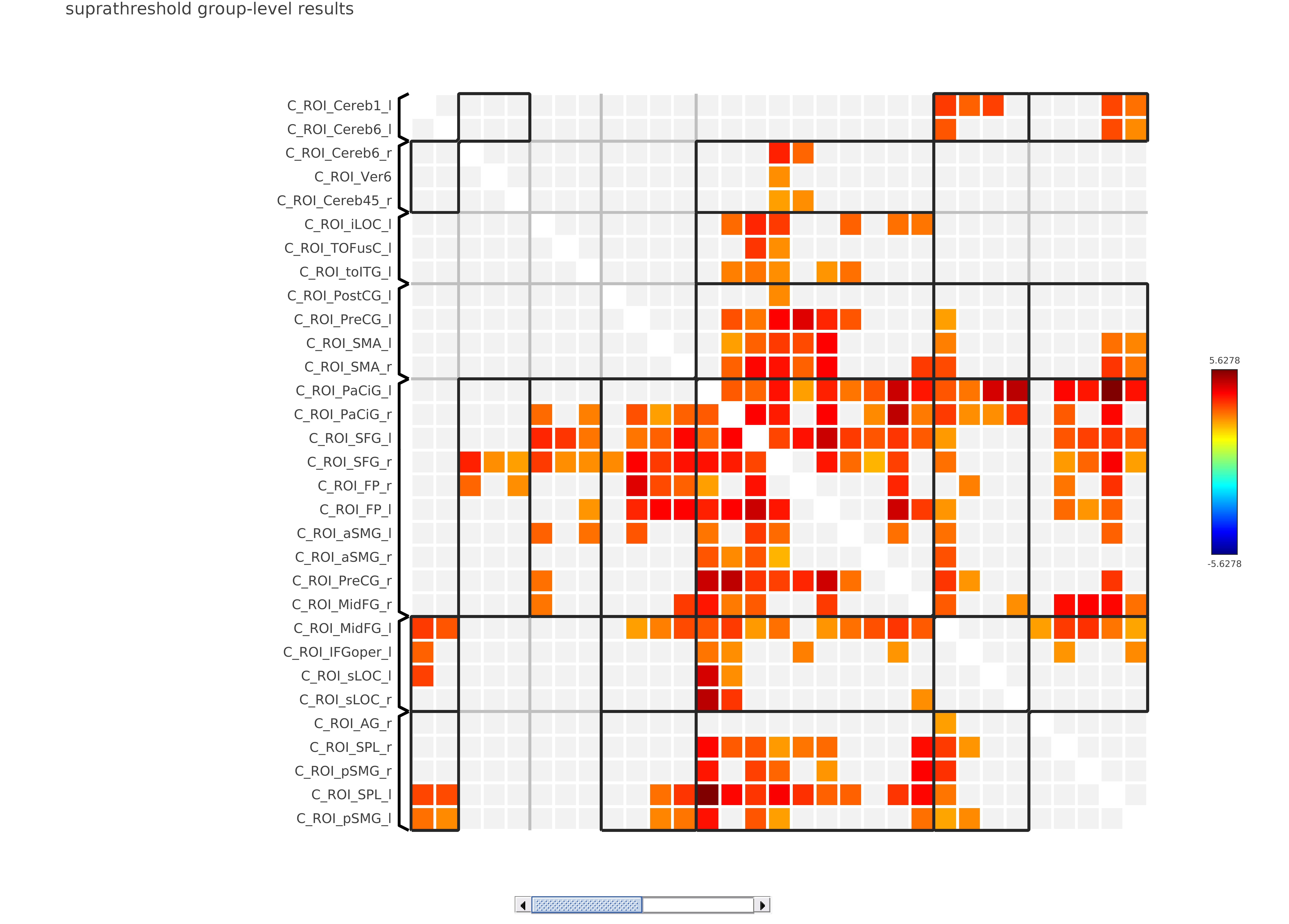

A. Dual Task

B. Single Cognitive Task

**Figure S2**. Group-level ROI-to-ROI functional connectivity results derived from cognitive-task ROIs under A: dual-task and B: single cognitive Go-task conditions. Each panel displays significant differences (*p* < 0.05, FDR-corrected) between older and younger adults (Old > Young) as a matrix plot, with color gradients representing t-values (warmer colors: stronger connectivity in older adults; cooler colors: stronger connectivity in younger adults).

**Figure S3.** Group-level ROI-to-ROI functional connectivity results derived from motor-task ROIs contrasting older and younger adults (Old > Young) under A: dual-task and B: single motor-task conditions. Each panel displays significant differences (*p* < 0.05, FDR-corrected) between older and younger adults (Old > Young) as a a matrix plot, with color gradients representing t-values (warmer colors: stronger connectivity in older adults; cooler colors: stronger connectivity in younger adults).

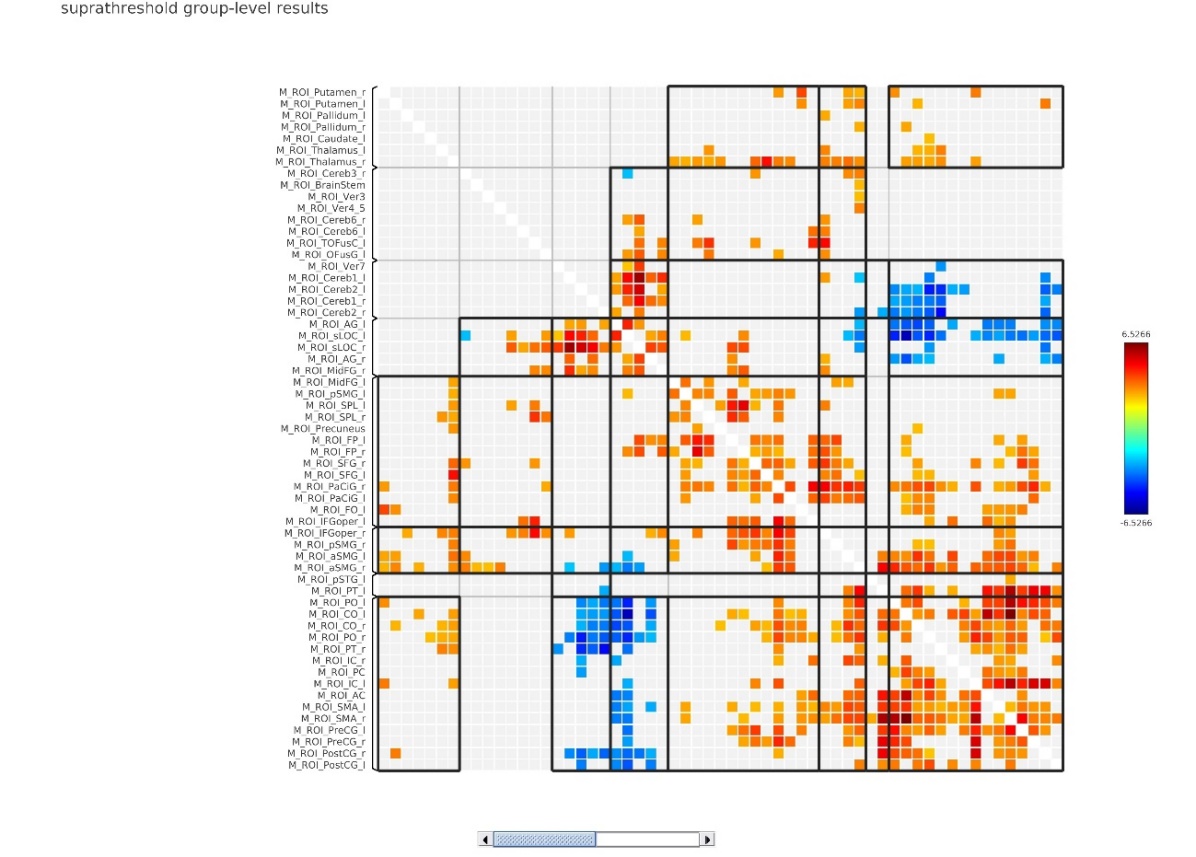

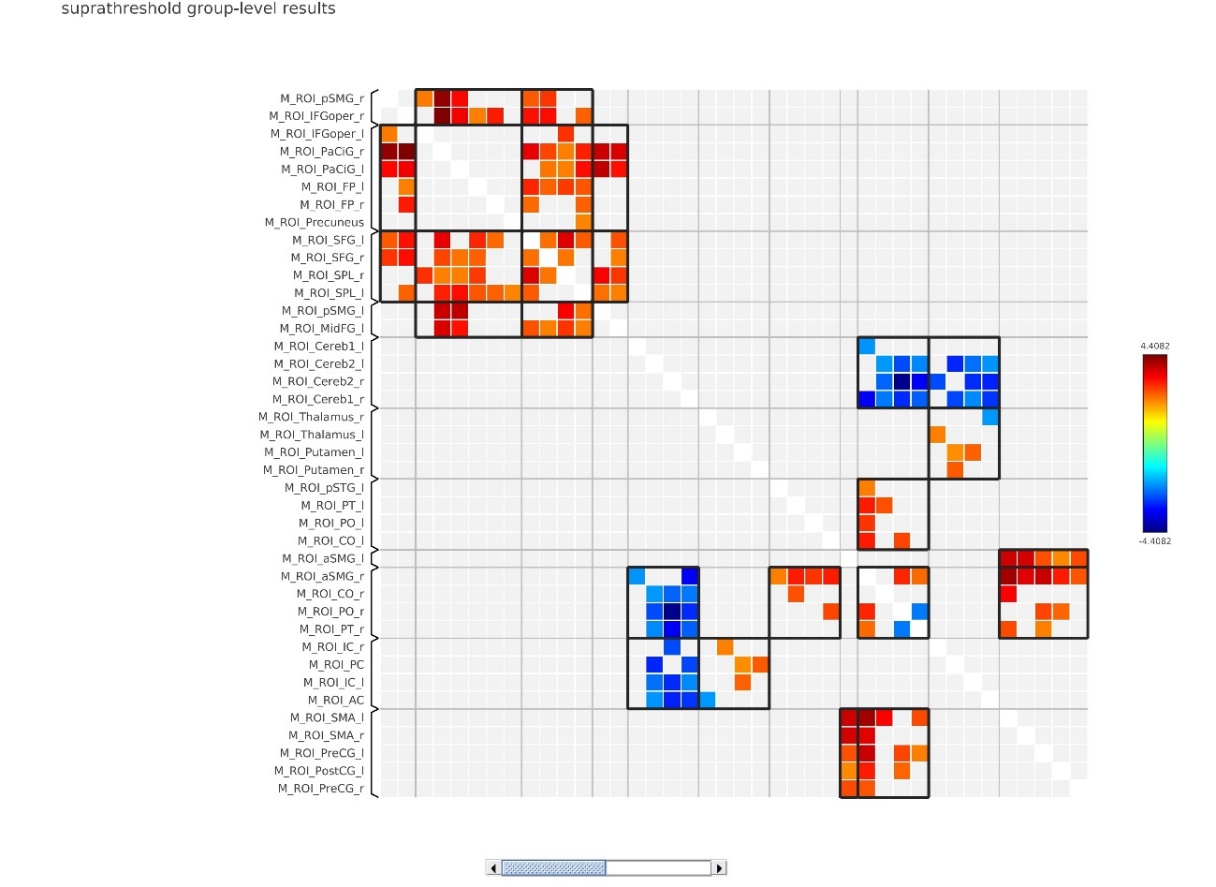

A. Dual Task

B. Single Motor Task

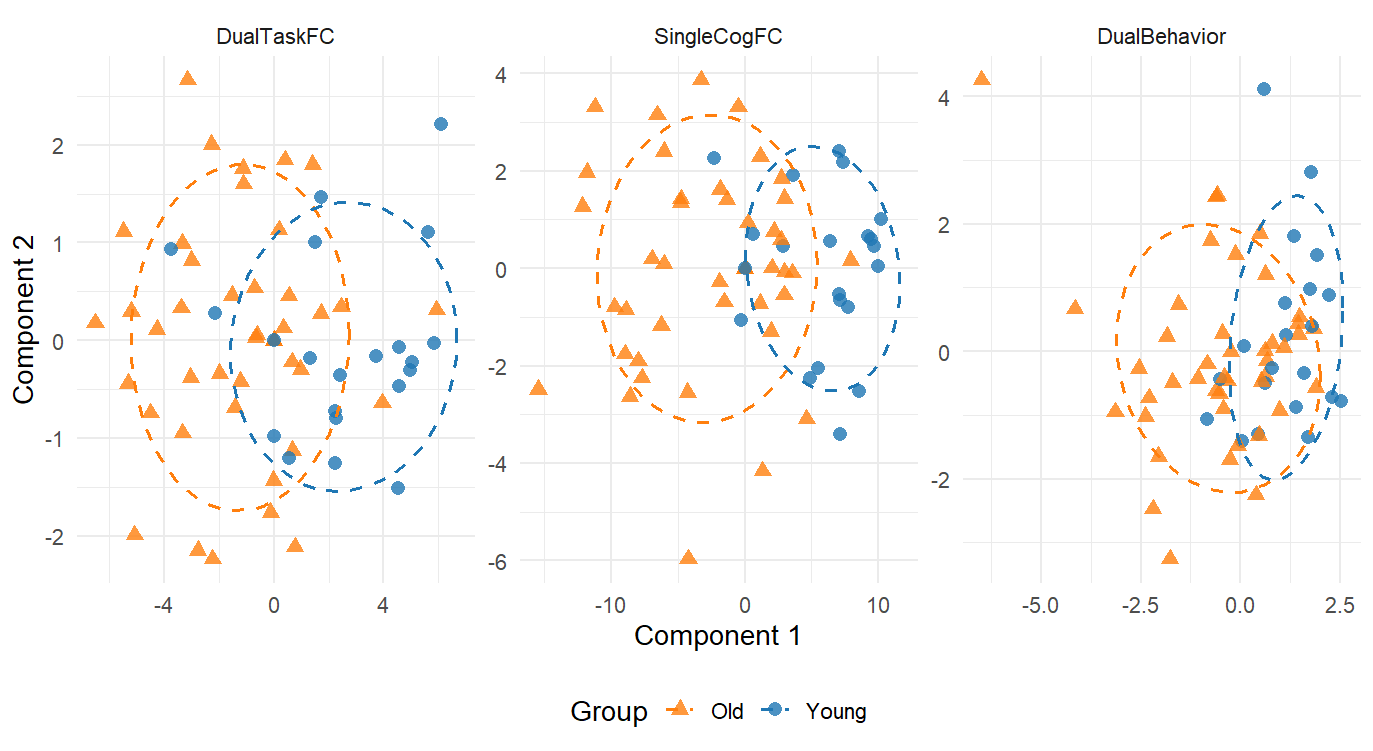

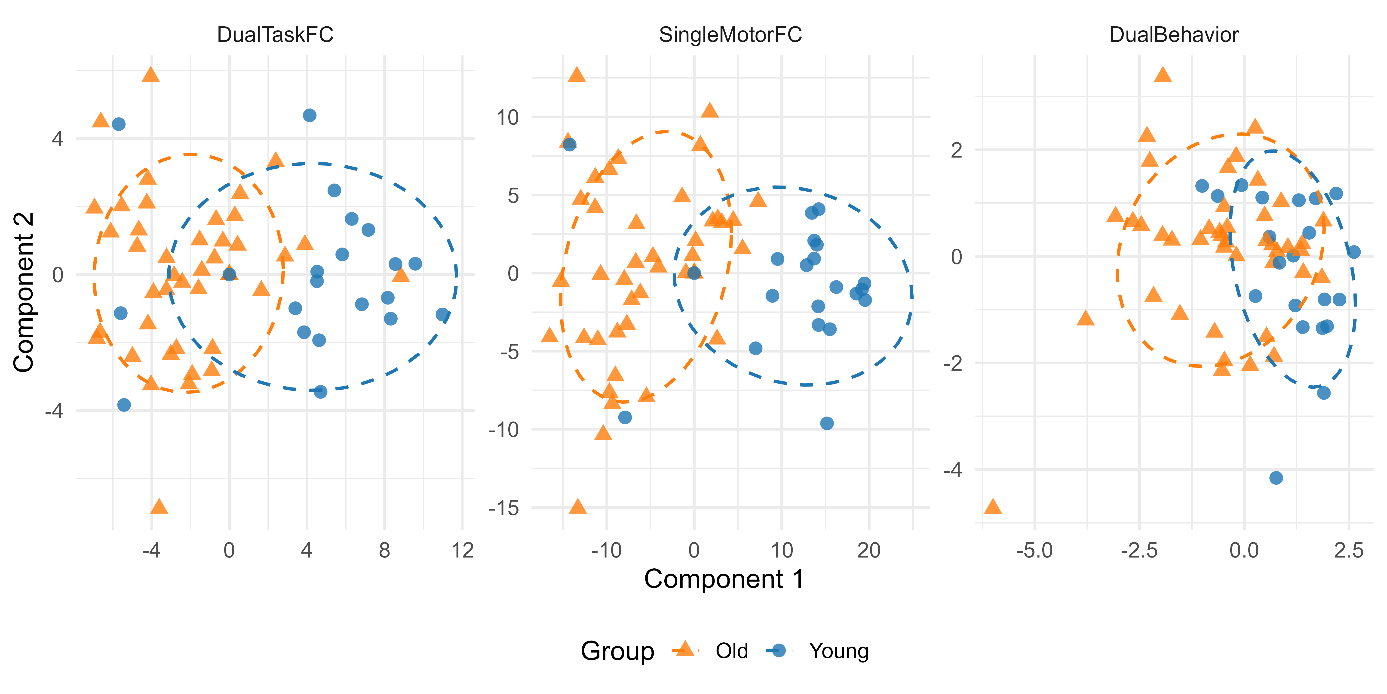

**Figure S4.** Multivariate component scores from RGCCA analysis plotted by task network and age group. The panel A displays component scores from the cognitive task network (Dual-Task FC, Single Cognitive FC, Dual Behaviour), while lower panel B shows corresponding scores from the Dual-Task FC, Single Motor FC, and Dual Behaviour blocks associated with the motor task network. Each plot depicts the scores of young (blue triangles) and older (red circles) adults on Component 1 and Component 2, with 68% confidence ellipses indicating group dispersion. The axis limits are tailored to the scale of each data block for better visual comparison. These plots illustrate distinct multivariate patterns in functional connectivity and behaviour across age groups and task demands. FC: functional connectivity.

##### References

Abraham, Alexandre, Fabian Pedregosa, Michael Eickenberg, Philippe Gervais, Andreas Mueller, Jean Kossaifi, Alexandre Gramfort, Bertrand Thirion, and Gael Varoquaux. 2014. “Machine Learning for Neuroimaging with Scikit-Learn.” Frontiers in Neuroinformatics 8. <https://doi.org/10.3389/fninf.2014.00014>.

Avants, B. B., C. L. Epstein, M. Grossman, and J. C. Gee. 2008. “Symmetric Diffeomorphic Image Registration with Cross-Correlation: Evaluating Automated Labeling of Elderly and Neurodegenerative Brain.” Medical Image Analysis 12 (1): 26–41. <https://doi.org/10.1016/j.media.2007.06.004>.

Behzadi, Yashar, Khaled Restom, Joy Liau, and Thomas T. Liu. 2007. “A Component Based Noise Correction Method (CompCor) for BOLD and Perfusion Based fMRI.” NeuroImage 37 (1): 90–101. <https://doi.org/10.1016/j.neuroimage.2007.04.042>.

Ciric, R., William H. Thompson, R. Lorenz, M. Goncalves, E. MacNicol, C. J. Markiewicz, Y. O. Halchenko, et al. 2022. “TemplateFlow: FAIR-Sharing of Multi-Scale, Multi-Species Brain Models.” Nature Methods 19: 1568–71. <https://doi.org/10.1038/s41592-022-01681-2>.

Dale, Anders M., Bruce Fischl, and Martin I. Sereno. 1999. “Cortical Surface-Based Analysis: I. Segmentation and Surface Reconstruction.” NeuroImage 9 (2): 179–94. <https://doi.org/10.1006/nimg.1998.0395>.

Esteban, Oscar, Ross Blair, Christopher J. Markiewicz, Shoshana L. Berleant, Craig Moodie, Feilong Ma, Ayse Ilkay Isik, et al. 2018. “fMRIPrep 24.1.1.” Software. <https://doi.org/10.5281/zenodo.852659>.

Esteban, Oscar, Christopher Markiewicz, Ross W Blair, Craig Moodie, Ayse Ilkay Isik, Asier Erramuzpe Aliaga, James Kent, et al. 2019. “fMRIPrep: A Robust Preprocessing Pipeline for Functional MRI.” Nature Methods 16: 111–16. <https://doi.org/10.1038/s41592-018-0235-4>.

Fonov, VS, AC Evans, RC McKinstry, CR Almli, and DL Collins. 2009. “Unbiased Nonlinear Average Age-Appropriate Brain Templates from Birth to Adulthood.” NeuroImage 47, Supplement 1: S102. <https://doi.org/10.1016/S1053-8119(09)70884-5>.

Gorgolewski, K., C. D. Burns, C. Madison, D. Clark, Y. O. Halchenko, M. L. Waskom, and S. Ghosh. 2011. “Nipype: A Flexible, Lightweight and Extensible Neuroimaging Data Processing Framework in Python.” Frontiers in Neuroinformatics 5: 13. <https://doi.org/10.3389/fninf.2011.00013>.

Gorgolewski, Krzysztof J., Oscar Esteban, Christopher J. Markiewicz, Erik Ziegler, David Gage Ellis, Michael Philipp Notter, Dorota Jarecka, et al. 2018. “Nipype.” Software. <https://doi.org/10.5281/zenodo.596855>.

Greve, Douglas N, and Bruce Fischl. 2009. “Accurate and Robust Brain Image Alignment Using Boundary-Based Registration.” NeuroImage 48 (1): 63–72. <https://doi.org/10.1016/j.neuroimage.2009.06.060>.

Jenkinson, Mark, Peter Bannister, Michael Brady, and Stephen Smith. 2002. “Improved Optimization for the Robust and Accurate Linear Registration and Motion Correction of Brain Images.” NeuroImage 17 (2): 825–41. <https://doi.org/10.1006/nimg.2002.1132>.

Klein, Arno, Satrajit S. Ghosh, Forrest S. Bao, Joachim Giard, Yrjö Häme, Eliezer Stavsky, Noah Lee, et al. 2017. “Mindboggling Morphometry of Human Brains.” PLOS Computational Biology 13 (2): e1005350. <https://doi.org/10.1371/journal.pcbi.1005350>.

Patriat, Rémi, Richard C. Reynolds, and Rasmus M. Birn. 2017. “An Improved Model of Motion-Related Signal Changes in fMRI.” NeuroImage 144, Part A (January): 74–82. <https://doi.org/10.1016/j.neuroimage.2016.08.051>.

Power, Jonathan D., Anish Mitra, Timothy O. Laumann, Abraham Z. Snyder, Bradley L. Schlaggar, and Steven E. Petersen. 2014. “Methods to Detect, Characterize, and Remove Motion Artifact in Resting State fMRI.” NeuroImage 84 (Supplement C): 320–41. <https://doi.org/10.1016/j.neuroimage.2013.08.048>.

Satterthwaite, Theodore D., Mark A. Elliott, Raphael T. Gerraty, Kosha Ruparel, James Loughead, Monica E. Calkins, Simon B. Eickhoff, et al. 2013. “An improved framework for confound regression and filtering for control of motion artifact in the preprocessing of resting-state functional connectivity data.” NeuroImage 64 (1): 240–56. <https://doi.org/10.1016/j.neuroimage.2012.08.052>.

Tustison, N. J., B. B. Avants, P. A. Cook, Y. Zheng, A. Egan, P. A. Yushkevich, and J. C. Gee. 2010. “N4ITK: Improved N3 Bias Correction.” IEEE Transactions on Medical Imaging 29 (6): 1310–20. <https://doi.org/10.1109/TMI.2010.2046908>.

Zhang, Y., M. Brady, and S. Smith. 2001. “Segmentation of Brain MR Images Through a Hidden Markov Random Field Model and the Expectation-Maximization Algorithm.” IEEE Transactions on Medical Imaging 20 (1): 45–57. <https://doi.org/10.1109/42.906424>.
